# GlnA3*_Mt_* is able to glutamylate spermine but it is not essential for the detoxification of spermine in *Mycobacterium tuberculosis*

**DOI:** 10.1101/2023.12.14.571729

**Authors:** Sergii Krysenko, Carine Sao Emani, Moritz Bäuerle, Maria Oswald, Andreas Kulik, Christian Meyners, Doris Hillemann, Matthias Merker, Inken Wohlers, Felix Hausch, Heike Brötz-Oesterhelt, Agnieszka Mitulski, Norbert Reiling, Wolfgang Wohlleben

**Affiliations:** Interfaculty Institute of Microbiology and Infection Medicine Tübingen (IMIT), Department of Microbiology and Biotechnology, Auf der Morgenstelle 28, University of Tübingen, Tübingen, Germany; Cluster of Excellence ’Controlling Microbes to Fight Infections’, University of Tübingen, Auf der Morgenstelle 28, 72076 Tübingen, Germany; Microbial Interface Biology, Research Center Borstel, Leibniz Lung Center, Parkallee 1-40, 23845 Borstel, Germany; Institute of Organic Chemistry and Biochemistry, Technical University Darmstadt, Darmstadt, Germany; National Reference Center for Mycobacteria, Research Center Borstel, Leibniz Lung Center, Parkallee 1-40, 23845 Borstel, Germany; Evolution of the Resistome, Research Center Borstel, Leibniz Lung Center, Parkallee 1-40, 23845 Borstel, Germany; Data Science, Research Center Borstel, Leibniz Lung Center, Parkallee 1-40, 23845 Borstel, Germany; Interfaculty Institute of Microbiology and Infection Medicine Tübingen (IMIT), Department of Microbial Bioactive Compounds, Auf der Morgenstelle 28, University of Tübingen, Tübingen, Germany; German Center for Infection Research (DZIF), Partner Site Hamburg-Lübeck-Borstel-Riems, Borstel, Germany; German Center for Infection Research (DZIF), Partner Site Tübingen, Tübingen, Germany; Centre for Synthetic Biology, Technical University of Darmstadt, 64283 Darmstadt, Germany; Present address: Valent BioSciences, 1910 Innovation Wy Suite 100, Libertyville, IL 60048, USA

**Keywords:** GS-like enzyme, GlnA3, glutamylation, polyamine metabolism, tuberculosis, infection, Rv3065

## Abstract

*Mycobacterium tuberculosis* is well adapted to survive and persist in the infected host, escaping the host immune response. Since polyamines, which are synthesized by infected macrophages are able to inhibit the growth of *M. tuberculosis*, the pathogen needs strategies to cope with toxic spermine. The actinomycete *Streptomyces coelicolor*, closely related to *M. tuberculosis* makes use of a gamma-glutamylation pathway to functionally neutralize spermine. We therefore considered whether a similar pathway would be functional in *M. tuberculosis*. In the current study we demonstrated that *M. tuberculosis* growth was inhibited by the polyamine spermine. Using a glutamine synthetase-based *in vitro* enzymatic activity assay we determined that GlnA3*_Mt_* (Rv1878) is a gamma-glutamylspermine synthetase. In an *in vitro* phosphate release assay we showed that purified His-Strep-GlnA3*_Mt_* as well as native GlnA3*_Mt_* prefer spermine as a substrate to putrescine, cadaverine, spermidine or other monoamines and amino acids, suggesting that GlnA3*_Mt_* may play a specific role in the detoxification of the polyamine spermine. However, the deletion of the *glnA3* gene in *M. tuberculosis* did not result in growth inhibition or enhanced sensitivity of *M. tuberculosis* in the presence of high spermine concentrations. Subsequent RNAsequencing of *M. tuberculosis* bacteria revealed that the gene cluster consisting of the efflux pump-encoding *rv3065-rv3066-rv3067* genes is upregulated upon spermine treatment, suggesting its involvement in bacterial survival under elevated spermine concentrations.

**IMPORTANCE:** Antibiotics for the treatment of *Mycobacterium tuberculosis* infections attack classical bacterial targets, such as the cell envelope or the ribosome. Upon *M. tuberculosis* infection macrophages synthesize the polyamine spermine which - at elevated concentrations - is toxic for *M. tuberculosis*. Based on our investigations of spermine resistance in the closely related actinomycete *Streptomyces coelicolor*, we hypothesized that the glutamyl-sperminesynthetase GlnA3 may be responsible for resistance against toxic spermine. Here we show that the mycobacterial glutamyl-sperminesynthetase indeed can inactivate spermine by glutamylation. However, GlnA3 is probably not the only resistance mechanism since a *glnA3* mutant of *M. tuberculosis* can survive under spermine stress. Gene expression studies suggest that an efflux pump may participate in resistance. The functional role of GlnA3*_Mt_* as well as of the spermine transporter in the pathogenicity of *M. tuberculosis* is of special interest for their validation as new targets of novel anti-tubercular drugs.

## 1. INTRODUCTION

Worldwide, tuberculosis (TB) remains the most prevalent, persisting and difficult to treat infectious disease and is associated with high mortality. In 2021, an estimated 10.6 million people developed TB disease, with an estimated 1.6 million deaths (1). Tuberculosis is caused by pathogenic mycobacteria of the *Mycobacterium tuberculosis* complex (MTBC), which are well adapted to survive and persist within infected patients (2). An overall increase in the global burden of multidrug-resistant (MDR) and rifampicin-resistant (RR) TB severely jeopardizes control of the TB epidemic as envisaged by the WHO End TB strategy. MDR/RR-TB has been identified on all continents with approximately 450.000 cases reported in 2021 (1). Treatment of MDR*-*TB infections is of particular difficulty due to the extended duration, poor safety, and high costs associated. Several antibiotics are effective in treating MDR-TB infections; however, these drugs often show limited efficacy and their use is coupled with diverse side-effects. Thus, the identification of pathways that are essential for mycobacterial growth *in vivo* would provide new targets for the rational design of more effective anti-TB agents that could be active against MDR-TB.

Polyamines are small aliphatic polyvalent cations, predominantly derived from amino acids such as ornithine, arginine, and lysine (3). They are widely distributed in nature and are present in all organisms, with the most common cellular polyamines being putrescine, cadaverine, spermidine and spermine. Polyamines have been implicated in a wide range of biological processes, and their intracellular levels are elevated predominantly during exposure to various stress conditions (3, 4). Thus, intracellular polyamine concentrations are tightly regulated by cellular metabolic pathways (4), as polyamine excess has been proven toxic for prokaryotic and eukaryotic organisms and can lead to cell death (5–8). Polyamines are able to interact with negatively charged molecules like RNA, DNA, proteins, polyphosphate, and phospholipids (9). Consequently, an imbalance in polyamine metabolism can significantly affect cellular homeostasis. An excess of polyamines can be detoxified by modifications such as glutamylation. Detoxification is also the first step in subsequent polyamine assimilation as C/N sources under nutrient limiting conditions. Polyamine catabolism was investigated in several bacterial species revealing that the polyamine utilization pathway is not universal for all bacteria. This process was studied extensively in Gram-negative bacteria such as *E. coli* and *P. aeruginosa* POA1. Polyamine utilization was reported to occur via the aminotransferase pathway (10), the γ-glutamylation pathway (10–12), the direct oxidation pathway for putrescine, and the spermine/spermidine dehydrogenase pathway. *E. coli* and *B. subtilis* were shown to acetylate spermidine (12–15), whereas in the actinobacterial model organism *Streptomyces coelicolor*, the γ-glutamylation pathway was demonstrated (16–18).

*S. coelicolor* was shown to synthesize two functional, glutamine synthetase-like (GS-like) enzymes for polyamine glutamylation: the gamma-glutamylpolyamine synthetases GlnA2*_Sc_* and GlnA3*_Sc_*. GlnA3*_Sc_* was reported to be a central enzyme for the γ-glutamylation pathway in *S. coelicolor*, and is highly specific for spermine (16, 17). In *S. coelicolor*, GlnA3*_Sc_*permits detoxification and subsequent utilization of polyamines as a nitrogen source and is indispensable for bacterial survival under high polyamine concentrations (16). Recent *in silico* analysis of *glnA*-like genes across the actinobacterial phyllum revealed numerous orthologues (*glnA2*, *glnA3* and *glnA4*) that code for proteins potentially involved in the colonization, persistence and survival of bacteria across diverse habitats (18).

Little is known about polyamine utilisation and its regulation in *M. tuberculosis.* In *M. tuberculosis*, GlnA3*_Mt_* has been annotated and classified as “GS-like” (19, 20). While the function, regulation and involvement of the glutamine synthetase (GS*_Mt_*) (GlnA1, Rv2220) in mycobacterial pathogenesis were extensively studied (19), the function of GlnA3*_Mt_* (Rv1878) so far remains unclear. Because many steps of nitrogen metabolism in *S. coelicolor* are nearly identical to those of *M. tuberculosis* (21–23), we hypothesized that *M. tuberculosis* might also exert a similar capacity for polyamine detoxification. The current study is the first to investigate polyamine metabolism in *M. tuberculosis* and the role of GlnA3*_Mt_* in this process.

## 2. RESULTS

### GlnA3*_Mt_* from *M. tuberculosis* restores the growth of the polyamine-deficient *S. coelicolor glnA3* mutant in polyamine-containing medium

As reported previously, GlnA3*_Sc_* confers the ability of *S. coelicolor* M145 to utilize polyamines as a sole nitrogen source (16). In order to test for a similar predicted function of the GlnA3*_Mt_*enzyme, the *glnA3_Mt_* gene was used for heterologous complementation of the *S. coelicolor* M145 Δ*glnA3_Sc_* mutant. The integrative vector pRM4 carrying a single copy of the *glnA3_Mt_* gene was integrated into the genome of the *S. coelicolor* M145 Δ*glnA3_Sc_*mutant. Successful integration of the construct in the *S. coelicolor* M145 genome was confirmed by PCR and sequencing. The phenotype of the complemented Δ*glnA3_Sc_* mutant was analyzed on complex, nitrogen-replete media (LB-agar, R5-agar), as well as on defined Evans-agar supplemented with single nitrogen sources (ammonium, L-glutamine or the polyamines putrescine, spermidine, spermine). Growth and morphology of the parental *S. coelicolor* M145 strain, the Δ*glnA3_Sc_*mutant and the Δ*glnA3_Sc_*complemented with *glnA3_Sc_* or *glnA3_Mt_* were monitored on solid media in a time range between 3 and 12 days of incubation at 30°C. The strains grew well on complex media (LB and R5) and on the defined Evans medium supplemented with ammonium and glutamine (Tab. 1).

**Tab. 1.** Physiological role of the *glnA3_Mt_* gene product in the *S. coelicolor glnA3* mutant grown in the presence of polyamines and other nitrogen sources.

| Medium | M145<br>(WT) | Δ*glnA3<sub>Sc</sub>* | Δ*glnA3<sub>Sc</sub>*<br>+*glnA3<sub>Mt</sub>* | Δ*glnA3<sub>Sc</sub>*<br>+*glnA3<sub>Sc</sub>* |
| --- | --- | --- | --- | --- |
| LB-Agar | +++ | +++ | +++ | +++ |
| R5-Agar | +++ | +++ | +++ | +++ |
| Evans-Agar<br>+ NH<sub>4</sub>Cl (50 mM) | +++ | +++ | +++ | +++ |
| Evans-Medium<br>+ Glutamine (50 mM) | +++ | +++ | +++ | +++ |
| Evans-Medium<br>+ Putrescine (200 mM) | +++ | - | + | ++ |
| Evans-Medium<br>+ Spermidine (25 mM) | +++ | - | + | ++ |
| Evans-Medium<br>+ Spermine (25 mM) | +++ | - | + | ++ |

Note: +: poor growth, ++: moderate growth, +++: strong growth; -: no growth.

The Δ*glnA3_Sc_* mutant was not able to grow on the defined Evans medium with polyamines as the only nitrogen source as previously reported (16). The heterologous complementation of the Δ*glnA3_Sc_* mutant by the *glnA3_Mt_* gene partially restored its growth on defined Evans medium supplemented with either putrescine, spermidine or spermine as the only nitrogen source (Tab. 1, Suppl. Fig. 1) thus demonstrating the functional equivalence of GlnA3*_Sc_* and GlnA3*_Mt_*. Interestingly, GlnA3*_Mt_* rescued growth of the *S. coelicolor* Δ*glnA3_Sc_* mutant grown on putrescine, although this enzyme has low substrate specificity towards short-chain polyamines (see Fig. 2). This indicates that the native GlnA2 enzyme, which is highly specific towards putrescine (16, 17), in the *S. coelicolor* test system could compensate for the lack of specificity towards putrescine by GlnA3*_Mt_*, allowing to rescue the growth of the mutant.

### GlnA3*_Mt_*catalyzes gamma-glutamylation of spermine and spermidine *in vitro*, but not of putrescine or cadaverine

To elucidate the biochemical function of GlnA3*_Mt_* and to confirm that GlnA3*_Mt_* is able to catalyze the predicted glutamylation reaction of organic polyamines, we applied a previously validated HPLC/mass spectrometry (HPLC/MS)-based assay (17). For this purpose, recombinant *glnA3_Mt_* was purified from *Escherichia coli* BL21 as a His-and Strep-tag fusion protein, with subsequent removal of the affinity tags. Analysis of the final product by size-exclusion chromatography supported the predicted dodecameric quaternary structure of GlnA3*_Mt_* (Fig. 1). The catalytic activity of purified GlnA3*_Mt_* was then tested by incubation with various polyamines in relevant reaction mixtures and the products analyzed using HPLC/MS analysis in negative MS mode (Fig. 1, reaction educts and products and their respective masses are depicted in Suppl. Fig. 2). Our data show that GlnA3*_Mt_* accepted glutamate and spermine as substrates in an ATP-dependent manner, resulting in the expected *m/z* for glutamylspermine. No glutamylated putrescine, cadaverine and spermidine were detected. The GlnA3*_Mt_*-catalyzed reaction generated a product with the mass-to-charge ratio of *m/z* 331, corresponding to the calculated mass of the gamma-glutamylspermine, supporting the hypothesis that GlnA3*_Mt_* functions as a gamma-glutamylspermine synthetase (Fig. 1).

**Fig. 1.**
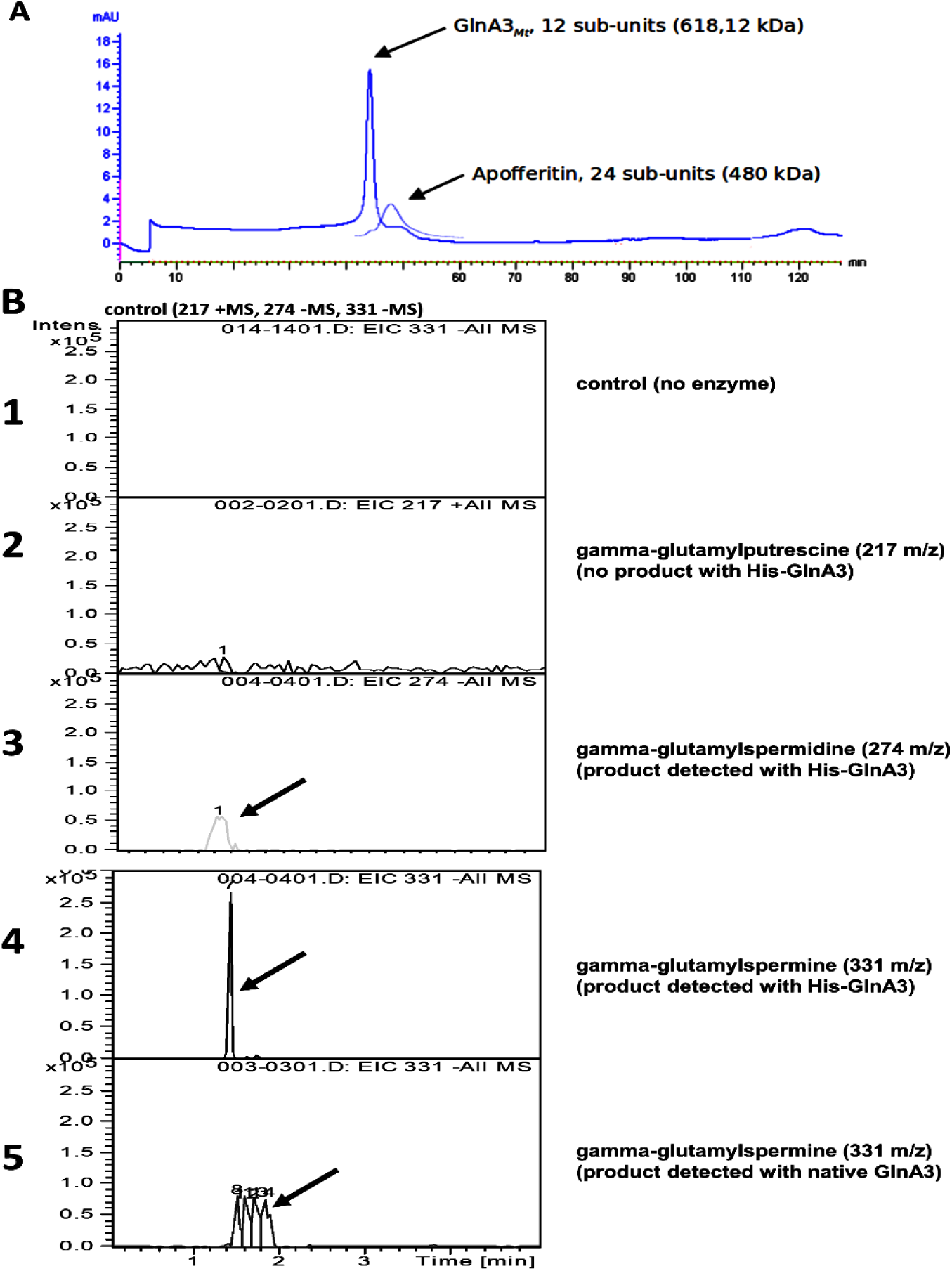
Purification of GlnA3Mt and generation of glutamylated spermine and spermidine by GlnA3Mt in an in vitro assay proved by the HPLC/ESI-MS analysis. A) purification of GlnA3Mt by size-exclusion chromatography (SEC). SEC supported the putative dodecameric quaternary structure of the enzyme. Pre-ordered apoferritin protein was used as a control for molecular mass estimation (elution profile shown in part). B) mass spectra chromatograms for the control detecting glutamylated putrescine, spermidine and spermine in a sample without GlnA3Mt enzyme (1), gamma-glutamylputrescine (m/z 217) (2), gamma-glutamylspermidine (m/z 274) (3), gamma-glutamylspermine (m/z 331), GlnA3Mt purified by SEC with the Tag (4), gamma-glutamylspermine (m/z 331), His-Strep-GlnA3Mt purified by affinity chromatography without the Tag (5). Glutamylated spermine and spermidine could be detected.

### Spermine is a preferred substrate for His-Strep-GlnA3*_Mt_*

To study the substrate specificity of GlnA3*_Mt_* and its kinetic parameters, an adapted GS activity assay (24) based on the detection of inorganic phosphate (Pi) (released from ATP hydrolysis) was employed. Recombinant His-Strep-GlnA3*_Mt_* or the *S. coelicolor* orthologue were incubated with test substrates (polyamines, monoamines, ammonium and amino acids) and required cofactors for 5 minutes, followed by quantification of inorganic phosphate release. All substrates were tested at 50 mM concentration as determined in a previous study in the *S. coelicolor* model system (16, 17). The results revealed that GlnA3*_Mt_* can accept polyamines and amino acids like methionine as substrates, but overall, it is very specific for spermine.

In order to compare the substrate specificity of GlnA3*_Mt_*with its homologue GlnA3*_Sc_*, which was characterized as a gamma-glutamylpolyamine synthetase in *S. coelicolor*, the adapted GS activity assay was applied as described above (Fig. 2).

**Fig. 2.**
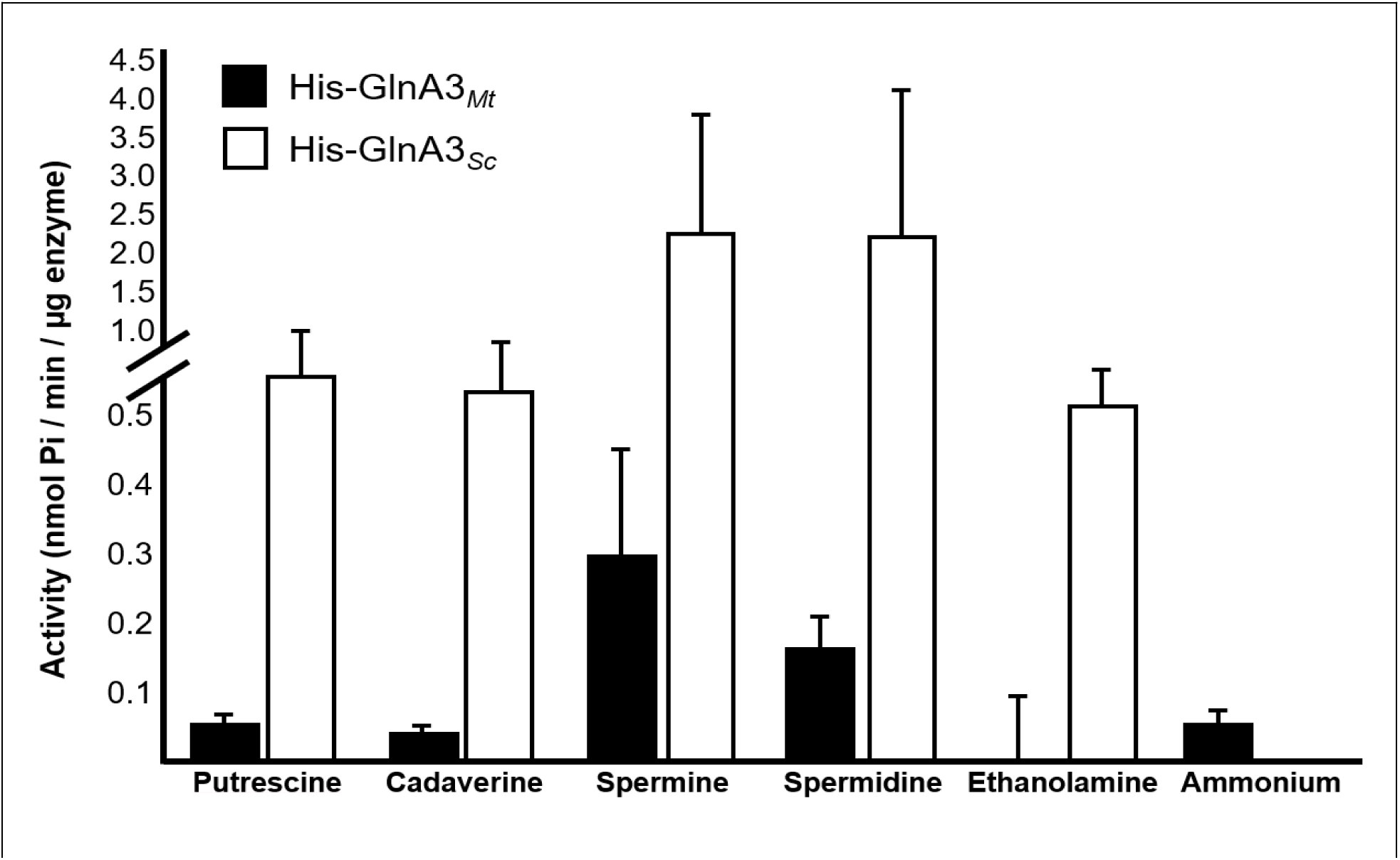
Specific activity of His-GlnA3*_Mt_* and His-GlnA3*_Sc_* with different nitrogen containing substrates. All substrates were at 50 mM. The mean value of n=3 biological replicates with n=3 technical replicates each with standard error is shown.

The analysis revealed that His-Strep-GlnA3*_Mt_* prefers spermine and spermidine as substrates. This finding is in contrast to His-GlnA3*_Sc_*which accepts also other polyamines (Fig. 2) and possesses a higher specific activity for all tested substrates except ammonium. This result demonstrated a rather specific spermine activity of GlnA3*_Mt_* from *M. tuberculosis*, which was not observed for its homolog GlnA3*_Sc_* from *S. coelicolor*.

In order to exclude putative effects of the tags on the obtained results, we independently assessed the activity of native GlnA3*_Mt_* with regard to the usage of putrescine, cadaverine, spermidine and spermine. The enzyme was purified as a fusion protein with the SUMO-Tag, which was subsequently removed by cleavage. The enzymatic activity of the native GlnA3*_Mt_* version was tested and compared with the His-Strep-GlnA3*_Mt_*. Although the native GlnA3*_Mt_* revealed overall higher enzymatic activity than the tagged version in *in vitro* assays, both GlnA3*_Mt_* variants showed significantly higher activity with spermine compared to other polyamine substrates (Suppl. Fig. 3). The effect remained the same after 1h of additional incubation.

### GlnA3*_Mt_*, a predicted gamma-glutamylpolyamine synthetase structurally resembles GlnA3*_Sc_*

GlnA3*_Mt_* shares 52% amino acid similarity across the full length sequence to GlnA3*_Sc_* from *S. coelicolor* M145 suggesting that the two orthologues may also share a similar tertiary structure (16).

In order to study the structural properties of GlnA3*_Mt_* in the absence of experimental data, a homology modelling approach was used. The cloud-based software SWISS-MODEL (25) was used to generate an *in silico* 3D model of the protein, based on the available amino acid sequence and a template model; in this case the previously solved structure of GlnA1*_Mt_* was used (26–30).

The GlnA3*_Mt_* model structure of *M. tuberculosis* comprises 12 identical subunits (Suppl. Fig. 4, A) organized in 2 rings, 6 sub-units in each (Suppl. Fig. 4, B), and one active site per sub-unit. Similar to both the solved GlnA*_Mt_*structure (27) and the GlnA3*_Sc_* model (16), a tunnel-like structure is observed in each active site of GlnA3*_Mt_*(Suppl. Fig. 4 B, C) comprising individual substrate binding sites for ATP, glutamate and ammonium. To improve confidence in the *in silico* data, we generated further models of GlnA3*_Mt_* using 20 different templates originating from various phyla of actinobacteria, proteobacteria, apicomplexa and cyanobacteria. Model quality was then assessed based on GMQE, QMEAN, and MolProbity scores (31), and Ramachandran analysis (Suppl. Fig. 4, D) (32, 33). Structure prediction was afterwards verified based on Alphafold predictions. The Alphafold prediction of the monomers matched the homology model monomer generated by the Chimera software.

The high diversity of templates used allowed the identification of conserved areas between the GlnA3*_Mt_* structure and other bacterial homologues. Across all analyzed proteins it was observed that most of the conserved amino acid residues were located in the active site of each monomer, namely in the glutamate, metal ions and ATP binding sites. The active site of GlnA3*_Mt_* contains 9 out of 12 conserved amino acid residues at the ammonium binding site compared to the described gamma-glutamylpolyamine synthetase GlnA3*_Sc_*, thought possessing more space on the loop for polyamine binding (no A169 residue in GlnA3*_Mt_*) (Fig. 3, Tab. 2).

**Tab 2.**
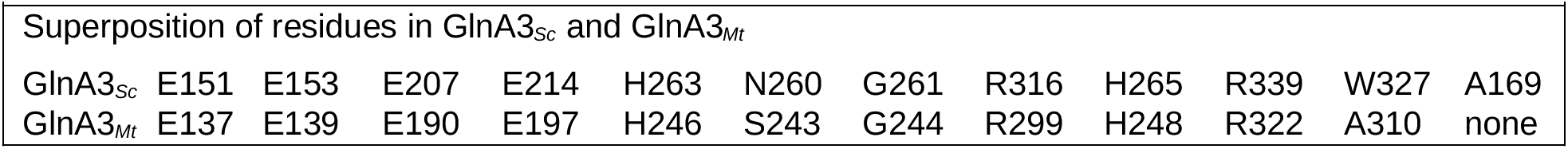

**Fig. 3.**
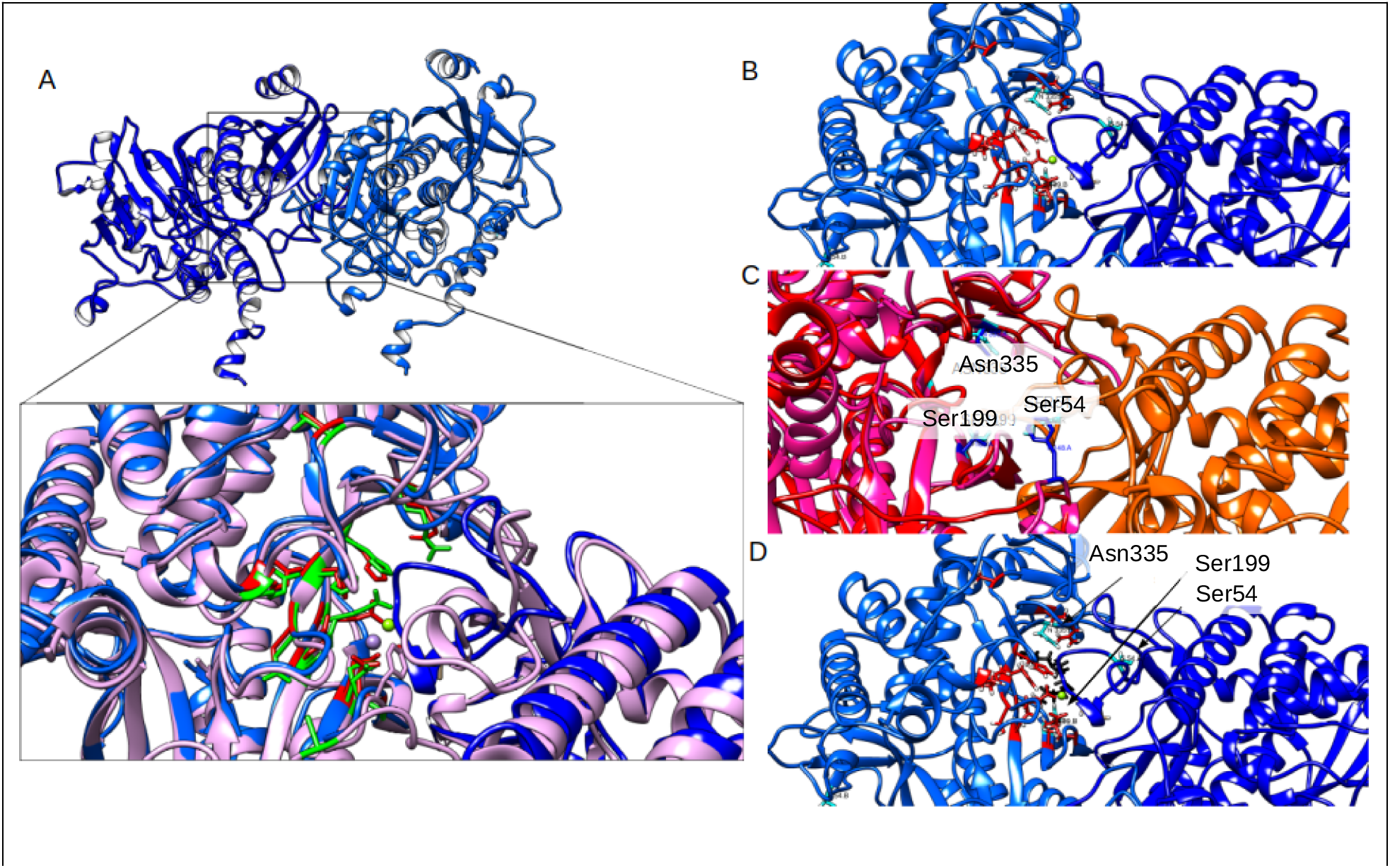
A) Structural alignment of the 3D model structure of the GlnA3*_Mt_* enzyme. Based on and superposed with the GlnA*_Mt_* template (PDB code 1BVC (27). Superposition of the GlnA3*_Sc_* (violet) and GlnA3*_Mt_* (blue) models with depicted key amino acids identified in GlnA3*_Sc_* (green, from (16) and GlnA3*_Mt_* (red) (see Tab. 2). B) Constellation of amino acid residues crucial for polyamine substrate binding in GlnA3*_Mt_.* The residues selected for site-directed mutagenesis i.e. Ser54, Ser199 and Asn335 (cyan) are depicted in the active site of GlnA3*_Mt_* among key residues in the binding pocket (red), C) computed localization of the spermine substrate (black) by molecular docking in the best scored dock position, D) structural alignment of structure models of two sub-units of GlnA3*_Mt_* (red and orange) overlaid with the crystal structure of PauA7*_Pa_* (violet).

### Site-directed mutagenesis of GlnA3*_Mt_* defines structural features important for spermine binding

In order to study the role of distinct residues of GlnA3*_Mt_* in catalytic activity and substrate recognition, a site-directed mutagenesis approach was undertaken. This approach relied on molecular docking with spermine and structural/sequence alignments with the available crystal structure of the gamma-glutamyl(mono-)polyamine synthetase from *P. aeruginosa* PauA7*_Pa_*(34), which is a functional homolog of GlnA3*_Mt_*. The comparison enabled the identification of potential key amino acids in the active site of GlnA3*_Mt_*. For example, Ser54 is part of the loop closing the active site, which is essential for substrate stabilization (34). In addition, Ser199 is a part of the so-called Tyr loop which partly forms the tunnel-like structure of the active site. Ser199 may also be part of the catalytic triad in GlnA3*_Mt,_* however this is yet to be defined. Finally, Asn335 forms part of a second loop structure that closes the active site (Fig. 3, A, C). The molecular modeling revealed that spermine may form interactions with those identified amino acid residues (Fig. 3, B). Based on these observations, these three amino acids (Ser54, Ser199, Asn335) were mutated via site directed mutagenesis and the respective recombinant proteins purified. Constructs containing the *glnA3_Mt_*\* variants containing a mutation were produced as previously described for the WT enzyme. Conducting activity and substrate specificity testing via the GS-based *in vitro* assay, it was demonstrated that the GlnA3*_Mt_*Ser199 enzyme variant exhibited no activity for any polyamine, in contrast to the WT and GlnA3*_Mt_*Ser54 and GlnA3*_Mt_*Asn335 variants (Fig. 4). These results were further verified by HPLC/MS analysis of reaction products (Suppl. Fig. 5).

**Fig. 4.**
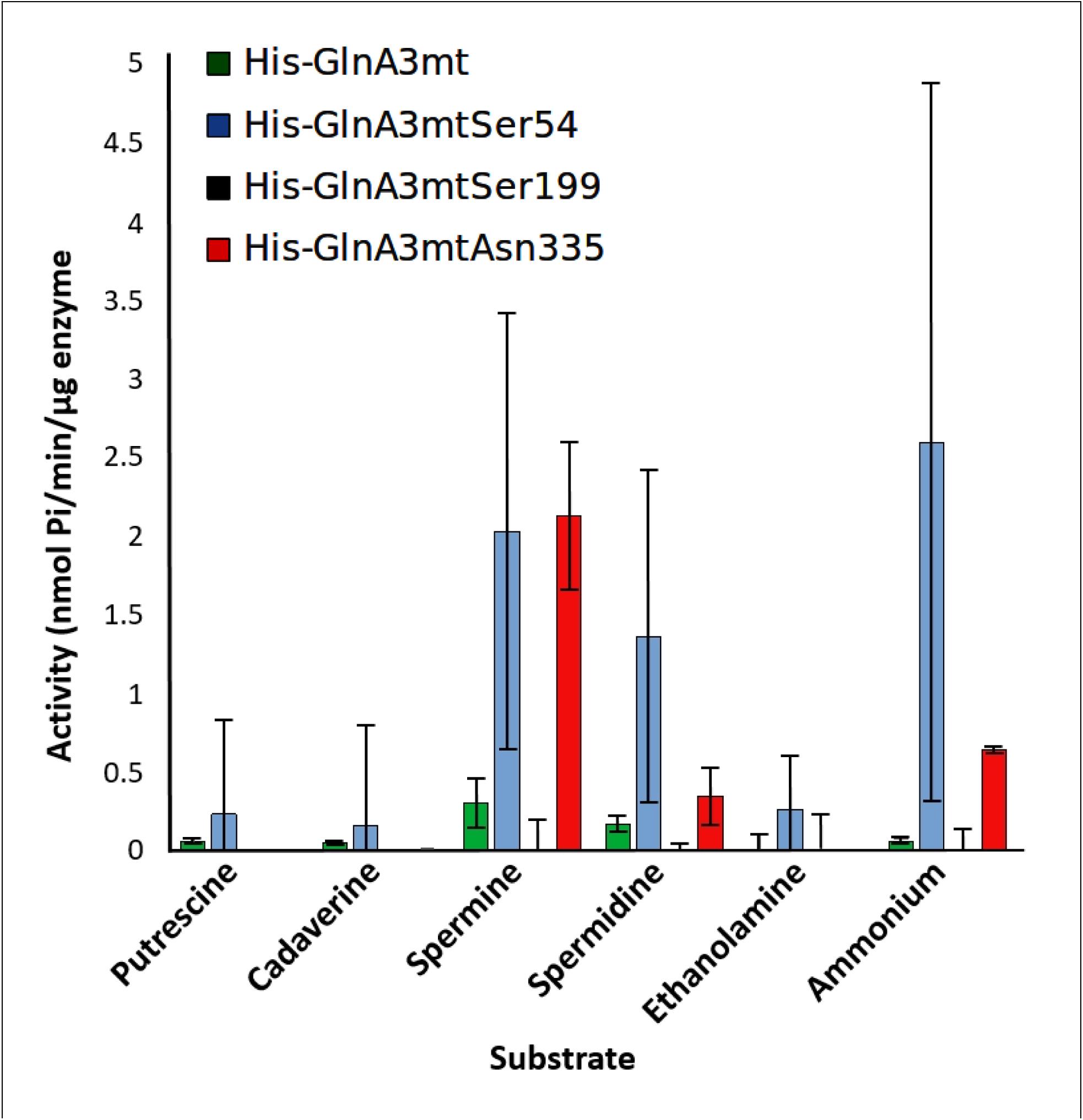
Specific activity of His-Strep-GlnA3*_Mt_* and His-Strep-GlnA3*_Mt_\** variants with different nitrogen-containing substrates. All substrates are at a concentration of 50 mM. The mean value of n=3 biological replicates with n=3 technical replicates each with standard error is shown.

### *M. tuberculosis* is able to detoxify the polyamine spermine

In order to assess potential toxicity of spermine against *M. tuberculosis*, we first investigated its effect on *M. tuberculosis* in the standard *M. tuberculosis* laboratory culture medium, 7H9, supplemented with albumin, dextrose and sodium chloride (ADS). Growth curve analyses revealed that 5 mM of spermine was required to achieve 90% inhibition (IC_90_) of *M. tuberculosis* growth (Fig. 5, A) with 50% inhibition (IC_50_) at 3 mM. Similar values were obtained irrespective of the supplement (ADS or OADC, see material and methods section) in a fluorescence-based test system employing an *M. tuberculosis* H37Rv strain expressing a red fluorescent protein (Suppl. Fig. 6, A and 6, B).

**Fig. 5.**
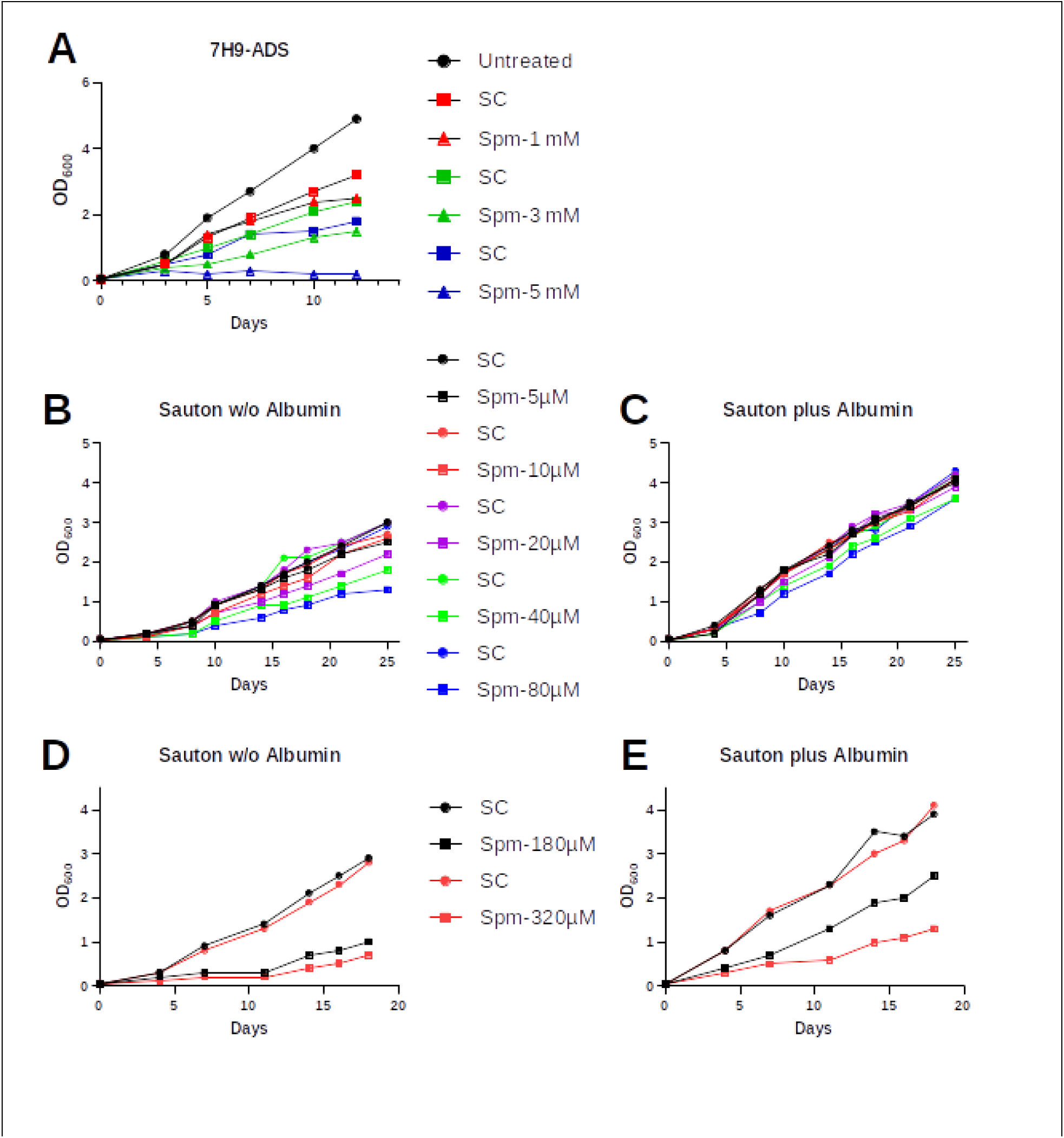
Growth of *M. tuberculosis* in the presence of spermine. Cells were incubated for a period of 14-26 days in **(A)** 7H9 media supplemented with ADS in the presence of 0.5-5 mM spermine (shown is one representative experiment out of four), **(B-E)** Sauton media in the presence of 5-80 µM (B, C) or 180-320 µM spermine (D, E) either incubated without (w/o) albumin (B, D) or with added albumin (C, E). SC: solvent control (DMSO) ; Spm: spermine

We then investigated the activity of spermine in a minimal medium without supplement (Sauton’s media), and observed that spermine was able to inhibit the growth of *M. tuberculosis* but at a substantially lower concentration (IC_90_ of 180-320 µM, IC_50_ of 80 µM (Fig. 5, B). Whenever albumin was added to the minimal media (3 mg/ml), the activity of spermine was inhibited (Fig. 5, C-E). This in accordance with previous studies demonstrating that spermine conjugates with albumin (35) thereby likely explaining the high MIC (low activity) of spermine in 7H9 standard media, containing albumin in the supplement.

In an additional we attempted to detect the glutamylated spermine in *M. tuberculosis,* the MTB Beijing strain (WT variant) and in the naturally occurring *glnA3* Beijing mutant were analysed by LC/MS. Low amounts of gamma-glutamylspermine were detected in these strains (Suppl. Fig. 7).

Since the Δ*glnA3* mutant was not more sensitive to Spermine relative to the wild-type, we referred to previous studies showing the implication of GlnA3_MT_ in nitrogen metabolism (19). Therefore, we investigated the ability of the mutant to use spermine as a sole C/N source. We found that the survival of the mutant was enhanced in the presence of spermine as the sole C/N source (Suppl. Fig. 8). This indicates that the mutant is able to metabolize spermine as a C/N source.

### The *M. tuberculosis ΔglnA3_Mt_* mutant is not sensitive to spermine

As GlnA3*_Sc_* plays a fundamental role in the survival of *S. coelicolor* at high spermine concentrations (16), we decided to employ a genetic approach to interrogate whether the same is true in *M. tuberculosis*. To this end, we constructed an unmarked, in-frame *glnA3_Mt_*-deletion mutant of *M. tuberculosis*, as described in Materials and Methods (36, 37). The *ΔglnA3_Mt_*(*Δrv1878*) mutant was identified by PCR, and validated by Southern blotting and targeted genome sequencing. The successful generation of the Δ*rv1878* mutant demonstrates that *rv1878* is not an essential gene in *M. tuberculosis* as previously indicated (38, 39).

The susceptibility of the mutant to spermine relative to the wild-type was first investigated by the broth microdilution method. The MIC was found to be similar to the wild-type’s. Next, spermine sensitivity was investigated by monitoring the growth of the strains in the presence of sub-inhibitory concentrations of spermine (half the MIC), Again, no difference in growth rate was observed between the mutant and wild-type (Fig 6, C and 6, D). Finally, in order to eliminate the possibility that the difference between the mutant and the wild-type was too small to detect with standard growth curve analyses or by MIC determinations, we evaluated the effect of spermine treatment on bacteria by determining colony-forming units (CFU). Three hours of treatment at excess concentrations of spermine (2 mM) in Sauton’s media revealed no difference in CFUs between the tested strains. In summary, all methods revealed that the mutant did not show an elevated sensitivity to spermine (Fig. 6, C, D, and E).

**Fig. 6.**
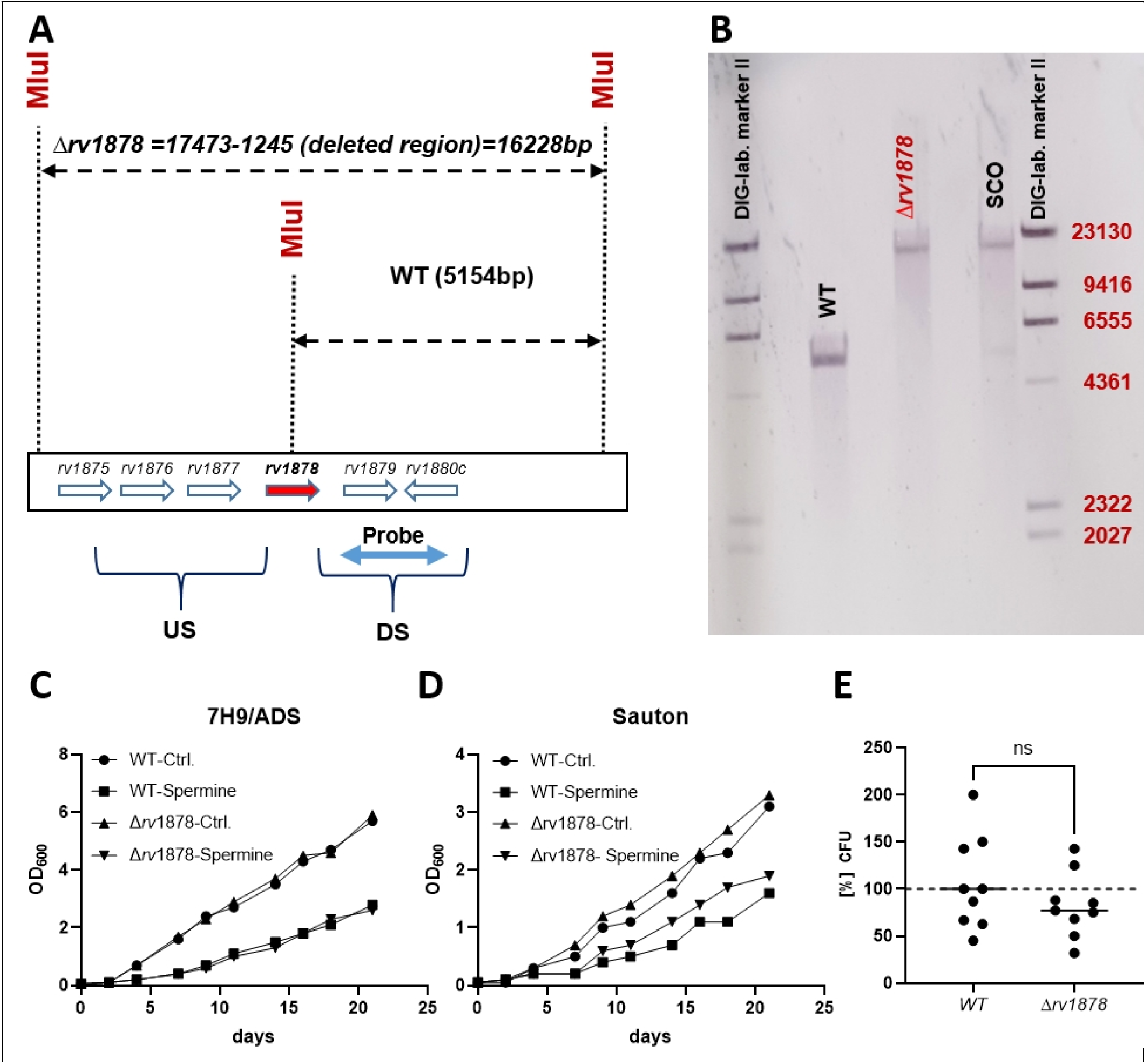
Generation and characterization of the *M. tuberculosis* Δ*glnA3* mutant relative to the wild-type. A and B) Southern blotting analyses to confirm gene deletion. The construct was designed such that the wild type DNA would yield a smaller band (5,154 bp) when digested with the enzyme *Mlu*I and hybridized with the probe flanking the DS (downstream) region, while the mutant DNA would yield a bigger band (16,228 bp) since *Mlu*I would not be able to cut in *glnA3*, since the restriction site has been deleted. The single cross-over mutant has both the wild-type and mutant bands. C) Sensitivity of the mutant relative to the wild type to 3 mM (1/2 MIC) spermine determined by growth curve analyses in 7H9 media, D) Sensitivity of the mutant relative to the wild-type to 80 µM (1/2 MIC) spermine determined by growth curve analyses in Sauton’s media, E) Sensitivity of the mutant relative to the wild type to 2 mM spermine determined by CFU counts and survival estimation (relative to the untreated control) in Sauton media for 3 hours). US, upstream; SCO Single cross over; ns, not significant)

### RNAseq analysis: Spermine does not regulate *Rv1878* and the adjacent mycobacterial genes

To investigate if *rv1878 (glnA3_Mt_)* and the adjacent genes (*rv1876*, *rv1877* and *rv1879*), were regulated by spermine stress, we initially performed RNAseq analysis of the wild-type bacteria cultured in 7H9 media and treated at half the MIC (3 mM), relative to the untreated control. A substantial change in gene expression of *rv1876*, *rv1878* and *rv1879* was not observed (less than 2fold (Fig. 7, A), validated by RT-PCR data (Fig. 7, B). Of note we observed in RNAseq analyses that *rv1877* displayed a 4-fold upregulation i.e. log2 median fold change of 1.7), which was shown to be even 10-fold enhanced, when validated by RT-PCR (Fig. 7, B) in 7H9 media conditions. Rv1877, a member of Major Facilitator Superfamily (MFS) has only recently been described as a multidrug efflux pump in *M. tuberculosis* (40). Since albumin could have possibly interfered with the activity of spermine (as discussed above) we repeated the experiment with medium lacking albumin to apply a more pronounced spermine stress. To this end RNA was sequenced from bacteria cultured in Sauton’s media, yet the regulation of all four genes found in this cluster was not altered by spermine stress (Fig. 7, C). These data show that *rv1877* is apparently upregulated only in supplemented 7H9 Media but not in Sauton’s media.

**Fig. 7.**
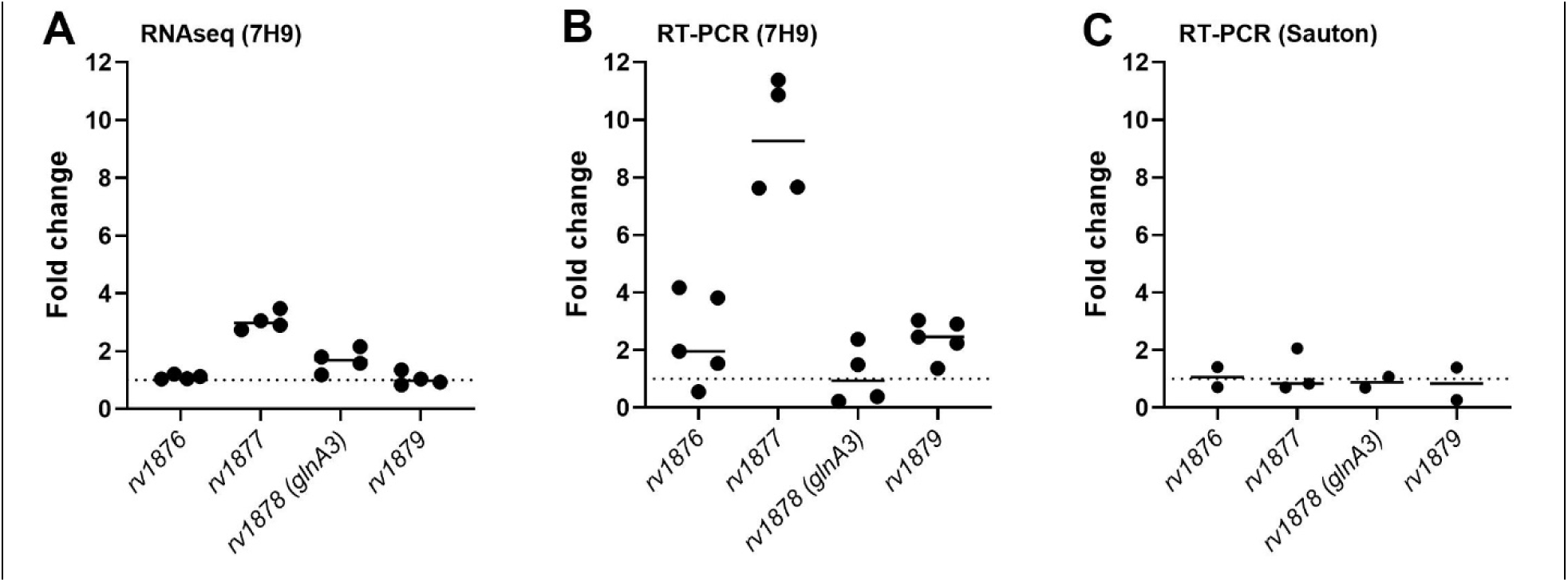
Alteration of the expression level of selected genes in *M. tuberculosis* WT. The expression was determined by RNAseq (A) and RT-PCR analysis (B, C) in 7H9 (A, B) and Sautońs (C) medium. Predicted gene functions: Rv1876 - bacterioferritin BfrA; Rv1877 - MFS transporter; Rv1878 – gamma-glutamylpolyamine synthetase GlnA3, Rv1879 - hypothetical protein

### RNAseq analysis: Spermine upregulates the gene coding for the efflux pump Rv3065

To investigate which other factors were possibly regulated by spermine stress, we compared the differences in gene expression of wild type *M. tuberculosis* with and without spermine treatment. For six genes with median raw read counts greater 20, a clearly reduced expression level with a log2 median fold change less than −3 was determined. In contrast, for 13 genes with sufficient number of reads the expression was increased by a log2 median fold change larger than 3 (Fig. 8).

**Fig. 8.**
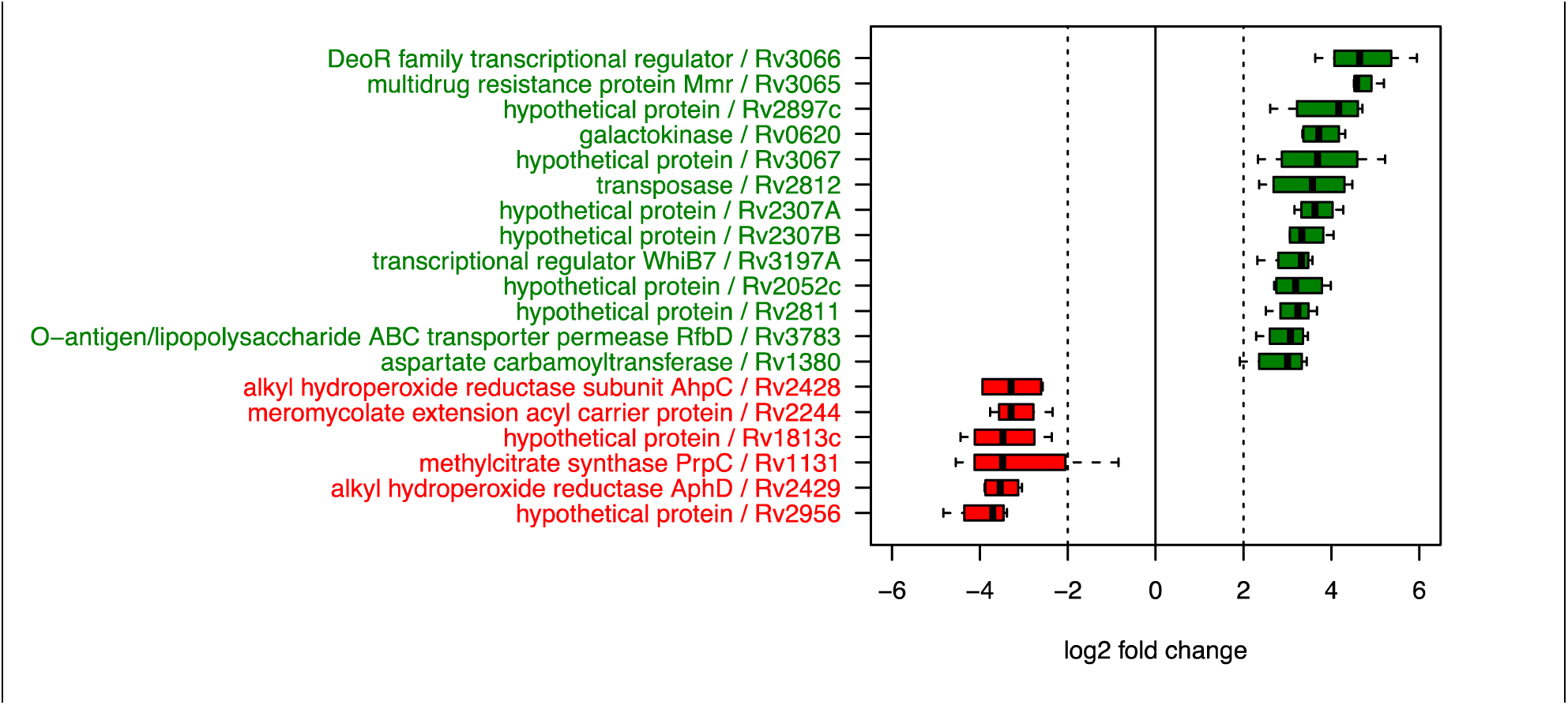
Expression of genes altered upon spermine treatment of *Mycobacterium tuberculosis* wild type in 7H9 media. Shown are box plots obtained from 4 independent experiments.

Among the five genes found most prominently upregulated were *rv0620* (log2 median fold change of 3.7; galactokinase) and *rv2897* (4.2 log2 fold; hypothetical protein), for which no specific role in the infection process or in polyamine metabolism is known. Interestingly, three other highly-upregulated genes (*rv3065, rv3066, rv3067*) are all located within the same genetic locus, probably forming an operon. The gene *rv3065* (log2 median fold change 4.6) codes for Mmr, a multidrug efflux pump that confers resistance to tetraphenylphosphonium (TPP), erythromycin, ethidium bromide, acriflavine, safranin O, pyronin Y and methyl viologen (41). The gene *rv3066* (log2 median fold change 4.6) codes for a putative transcriptional regulator of the DeoR family, while *rv3067* (log2 median fold change 3.7) encodes an uncharacterized protein, localised in the plasma membrane (https://www.uniprot.org/uniprotkb/I6YB21/entry). Bioinformatic analyses and previous studies (41) revealed that these proteins are neither found in *M. smegmatis* nor in *S. coelicolor.* Thus, it is tempting to speculate that Rv3065 is an important efflux pump, responsible for the excretion of polyamines and thus enabling *M. tuberculosis* to survive in excess spermine. Furthermore, in the model system *S. coelicolor*, a full transcriptome RNAseq analysis was performed in the presence of polyamines putrescine, cadaverine, spermidine and spermine in the wild type and in the *glnA3* mutant (17). This study supported our RNAseq results obtained in this work, and in connection with other experiments performed provided evidence that the second GlnA-like enzyme GlnA2 is involved in polyamine glutamylation. However, further investigation of this hypothesis was beyond the scope of this study.

## 3. MATERIALS AND METHODS

### Strains and growth conditions

*M. tuberculosis* (MTB) H37Rv was obtained from ATCC (27294) and genetically manipulated to obtain the mutant. The fluorescence expressing strain is an H37Rv strain carrying a shuttle vector (pCherry) (42, 43) expressing (under P_smyc_) *m*Cherry (a variant of the *Discosoma sp* red fluorescent protein (D*s*Red) (44). Strains were cultured either in Difco Middlebrook 7H9 broth (Becton Dickinson, Heidelberg, Germany) supplemented with ADS (0.005 g/ml Bovine albumin fraction V, 0.002 g/ml dextrose, 0.85 mg/ml sodium chloride) containing 0.05% tyloxapol (Sigma Aldrich), 0.02% (v/v) glycerol, or OADC (0.05 mg/ml oleic acid, 0.005 g/ml Bovine albumin fraction V, 0.002 g/ml dextrose, 0.004 mg/ml catalase, 0.85 mg/ml sodium chloride), or cultured in the ready-made Sauton’s media (HIMEDIA) containing glycerol (0.02%) and tyloxapol (0.05%). The solid media used for plating was 7H11 agar based (Sigma Aldrich) supplemented with either ADS or OADC. Spermine was obtained from Sigma Aldrich, and 100 mM stocks were made in DMSO and stored at −20°C for downstream experiments.

The parental strain *S. coelicolor* M145 as well as all mutants were incubated for 4-5 days at 30°C on the defined Evans-agar base (modified after (45) supplemented with following nitrogen sources: 50 mM ammonium chloride, 50 mM L-glutamine, 200 mM putrescine dihydrochloride, 25 mM spermidine trihydrochloride, and 25 mM spermine tetrahydrochloride in appropriate concentrations. Genetic manipulation of *S. coelicolor* M145 was performed as described previously (46, 47).

### Growth curve and susceptibility testing of *M. tuberculosis*

Frozen stocks were used to inoculate starter cultures, after 7-10 days of incubation, these were used to sub-culture in various growth conditions, 7H9-ADS or Sauton’s media, with and without spermine. For each tested concentration of spermine, a corresponding, untreated, DMSO control was included. The OD_600_ was measured and recorded every second day. To independently validate the data by fluorescence measurements *M. tuberculosis* growth was monitored using an mCherry expressing *M. tuberculosis* H37Rv strain which was analyzed at various time points, in a 96-well dark plate, using a multimode plate reader (Synergy 2, Biotek/Agilent) (55). The minimum inhibitory concentration (MIC) was also determined with the broth-microdilution method, using resazurin to enable colorful detection of growth as previously described (56, 57). In order to evaluate the susceptibility of the strains by CFUs count, logarithmic phase cultures, were adjusted to an OD_600_ of 0.2, this then was diluted 100X, to have approximately 10^4^ CFUs/ml, and this was then exposed to either 2 mM spermine or the DMSO-control for about 3 hours in a 96-well plate, at 37°C. These were serially diluted and plated on solid media that were incubated for up to 3 weeks. The CFUs of the treated culture was divided by the CFUs of the untreated DMSO control culture, in order to determine the percentage survival, this was repeated at least 8 times.

### Generation and complementation of the *M. tuberculosis* Δ*glnA3* mutant

This was achieved as previously described (36, 37). Briefly, approximately 2500 bp fragments upstream (US) and downstream (DS) *rv1878* (*glnA3*) were amplified using pfu high fidelity GC rich target polymerase (Agilent) primers listed in Table 3. These were subsequently cloned into the pJET sub-clonig vector (CloneJET PCR Cloning Kit, Thermofischer). Then the US and DS fragments were restricted using enzymes *Bsr*GI, S*pe*I and *Hin*dIII, which were incorporated in the primer design (see Table 3). Using a two-way ligation method, the restricted US and DS fragments were ligated to the p2NIL suicide plasmid (36), previously linearized with *Bsr*GI and *Hin*dIII. Then the *Pac*I restriction fragment containing, *lacZ*, and *sacB* genes was restricted from pGOAL17 (36) and cloned into the *Pac*I site of p2NIL. The phage resistant and endonuclease I (*endA1*) deficient *E. coli* strain D10-beta (New England Biolabs Inc.) was used in the entire cloning process. The integrity of the inserts was verified by Sanger sequencing at the end of each cloning step, throughout the cloning process (sequencing primers found in Table 3). The resulting construct p2NIL-A3-US/DS-G17, was used to transform *M. tuberculosis*, and the mutant was generated through two subsequent homologous recombination events as previously described (36). The mutant was screened by PCR (primers in Table 3), confirmed by Southern blotting as previously described (37) and by targeted genome sequencing (58). The mutant was complemented as previously described (37), using the pMVhsp60 integrative vector (59), where *glnA3* was cloned downstream the heat shock promoter (P_hsp60_), to maintain constitutive expression at 37°C (Table 3).

**Tab. 3.**
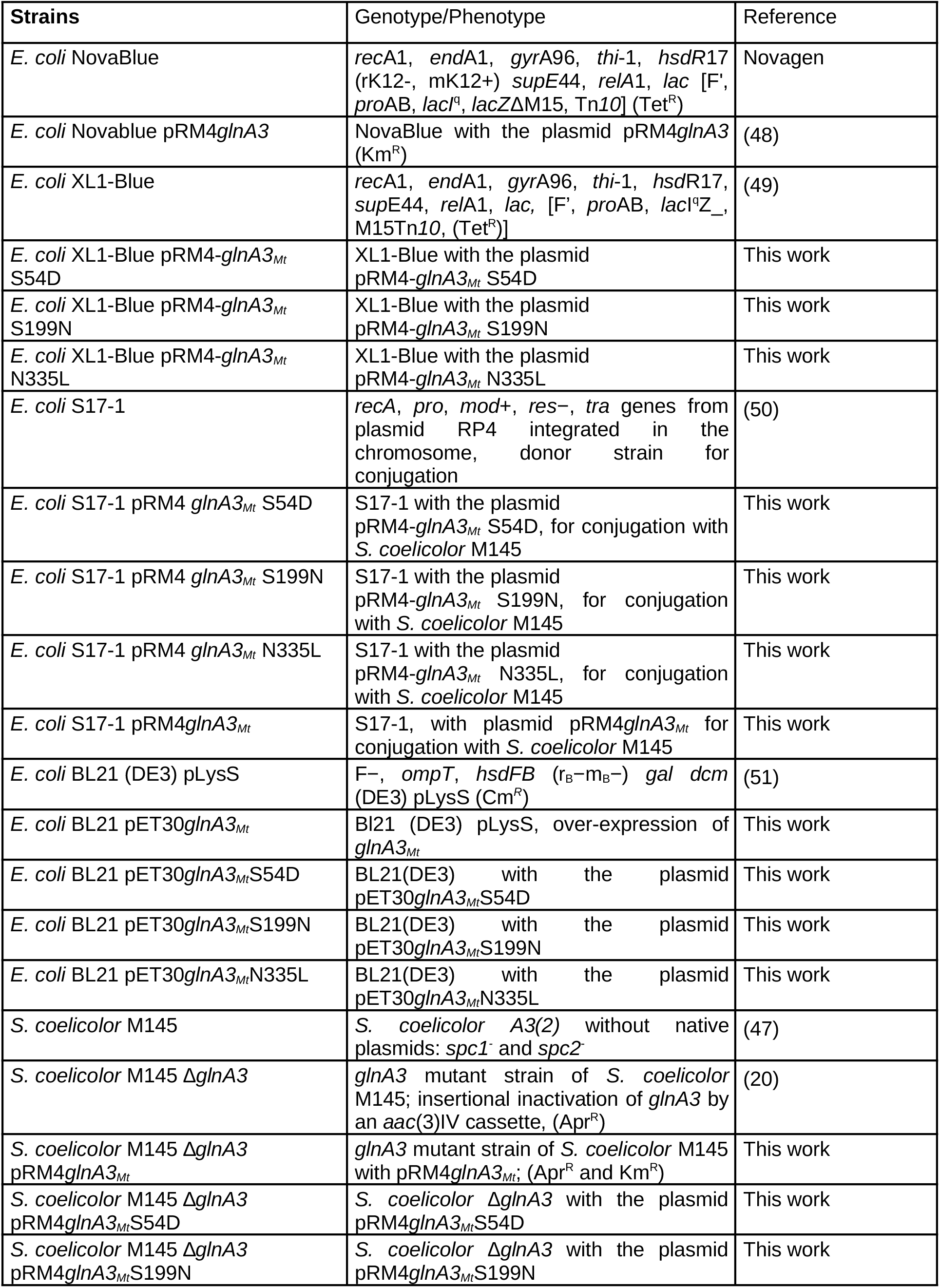
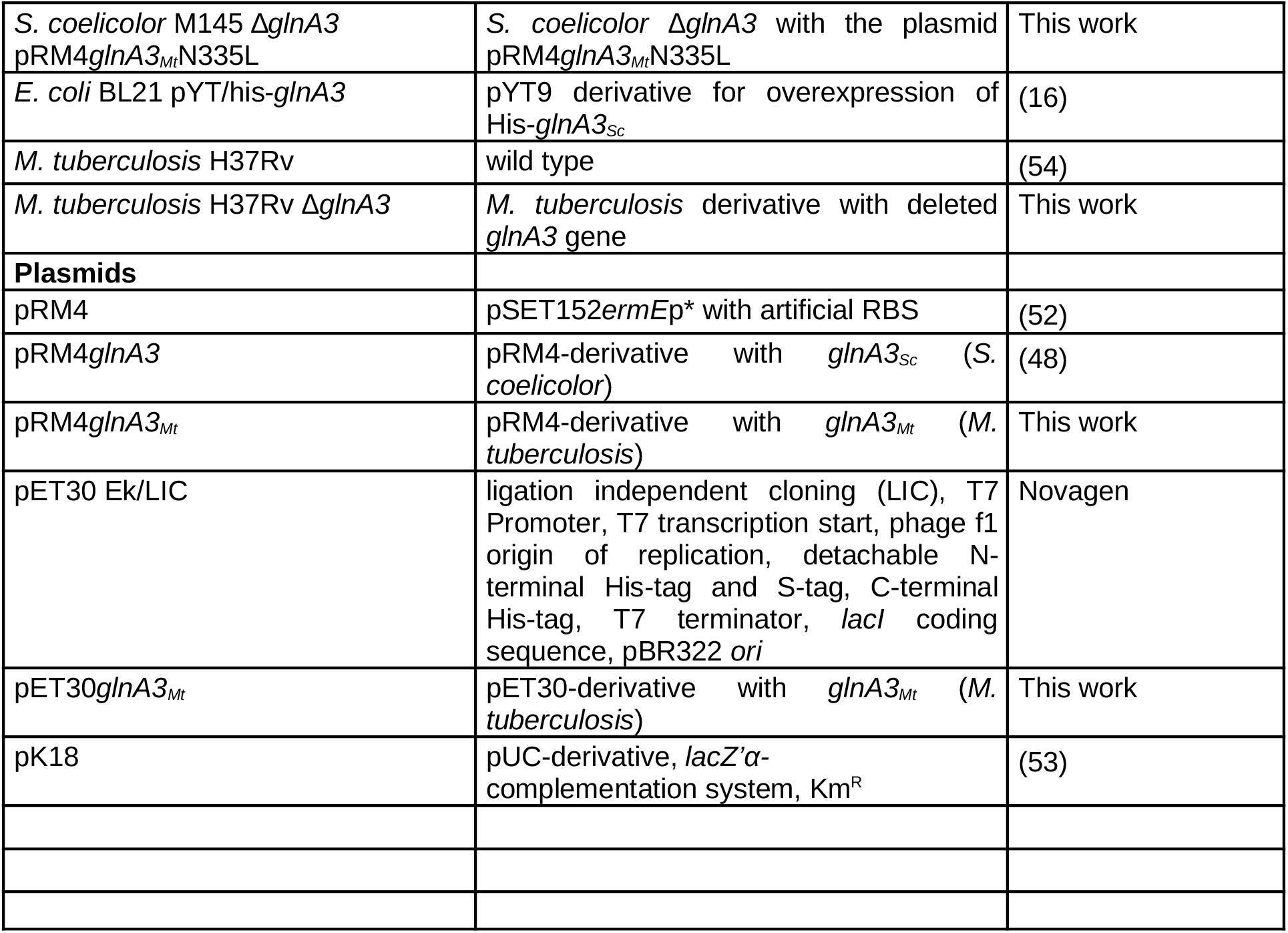
Strains and plasmids used in this study.

| Strains | Genotype/Phenotype | Reference |
| --- | --- | --- |
| *E. coli* NovaBlue | *recA1, endA1, gyrA96, thi-1, hsdR17* (rK12-, mK12+) *supE44, relA1, lac* [F', *proAB, lacI<sup>q</sup>, lacZΔM15, Tn10*] (Tet<sup>R</sup>) | Novagen |
| *E. coli* Novablue pRM4*glnA3* | NovaBlue with the plasmid pRM4*glnA3* (Km<sup>R</sup>) | (48) |
| *E. coli* XL1-Blue | *recA1, endA1, gyrA96, thi-1, hsdR17, supE44, relA1, lac*, [F', *proAB, lacI<sup>q</sup>Z*_, M15Tn10, (Tet<sup>R</sup>)] | (49) |
| *E. coli* XL1-Blue pRM4-*glnA3*<sub>Mt</sub> S54D | XL1-Blue with the plasmid pRM4-*glnA3*<sub>Mt</sub> S54D | This work |
| *E. coli* XL1-Blue pRM4-*glnA3*<sub>Mt</sub> S199N | XL1-Blue with the plasmid pRM4-*glnA3*<sub>Mt</sub> S199N | This work |
| *E. coli* XL1-Blue pRM4-*glnA3*<sub>Mt</sub> N335L | XL1-Blue with the plasmid pRM4-*glnA3*<sub>Mt</sub> N335L | This work |
| *E. coli* S17-1 | *recA, pro, mod+*, *res-*, *tra* genes from plasmid RP4 integrated in the chromosome, donor strain for conjugation | (50) |
| *E. coli* S17-1 pRM4 *glnA3*<sub>Mt</sub> S54D | S17-1 with the plasmid pRM4-*glnA3*<sub>Mt</sub> S54D, for conjugation with *S. coelicolor* M145 | This work |
| *E. coli* S17-1 pRM4 *glnA3*<sub>Mt</sub> S199N | S17-1 with the plasmid pRM4-*glnA3*<sub>Mt</sub> S199N, for conjugation with *S. coelicolor* M145 | This work |
| *E. coli* S17-1 pRM4 *glnA3*<sub>Mt</sub> N335L | S17-1 with the plasmid pRM4-*glnA3*<sub>Mt</sub> N335L, for conjugation with *S. coelicolor* M145 | This work |
| *E. coli* S17-1 pRM4*glnA3*<sub>Mt</sub> | S17-1, with plasmid pRM4*glnA3*<sub>Mt</sub> for conjugation with *S. coelicolor* M145 | This work |
| *E. coli* BL21 (DE3) pLysS | F<sup>-</sup>, *ompT, hsdFB* (r<sub>B</sub>-m<sub>B</sub>-) *gal dcm* (DE3) pLysS (Cm<sup>R</sup>) | (51) |
| *E. coli* BL21 pET30*glnA3*<sub>Mt</sub> | BL21 (DE3) pLysS, over-expression of *glnA3*<sub>Mt</sub> | This work |
| *E. coli* BL21 pET30*glnA3*<sub>Mt</sub>S54D | BL21(DE3) with the plasmid pET30*glnA3*<sub>Mt</sub>S54D | This work |
| *E. coli* BL21 pET30*glnA3*<sub>Mt</sub>S199N | BL21(DE3) with the plasmid pET30*glnA3*<sub>Mt</sub>S199N | This work |
| *E. coli* BL21 pET30*glnA3*<sub>Mt</sub>N335L | BL21(DE3) with the plasmid pET30*glnA3*<sub>Mt</sub>N335L | This work |
| *S. coelicolor* M145 | *S. coelicolor* A3(2) without native plasmids: *spc1*<sup>-</sup> and *spc2*<sup>-</sup> | (47) |
| *S. coelicolor* M145 Δ*glnA3* | *glnA3* mutant strain of *S. coelicolor* M145; insertional inactivation of *glnA3* by an *aac(3)IV* cassette, (Apr<sup>R</sup>) | (20) |
| *S. coelicolor* M145 Δ*glnA3* pRM4*glnA3*<sub>Mt</sub> | *glnA3* mutant strain of *S. coelicolor* M145 with pRM4*glnA3*<sub>Mt</sub>; (Apr<sup>R</sup> and Km<sup>R</sup>) | This work |
| *S. coelicolor* M145 Δ*glnA3* pRM4*glnA3*<sub>Mt</sub>S54D | *S. coelicolor* Δ*glnA3* with the plasmid pRM4*glnA3*<sub>Mt</sub>S54D | This work |
| *S. coelicolor* M145 Δ*glnA3* pRM4*glnA3*<sub>Mt</sub>S199N | *S. coelicolor* Δ*glnA3* with the plasmid pRM4*glnA3*<sub>Mt</sub>S199N | This work |
| *S. coelicolor* M145 $\Delta glnA3$ pRM4*glnA3*<sub>Mt</sub>N335L | *S. coelicolor* $\Delta glnA3$ with the plasmid pRM4*glnA3*<sub>Mt</sub>N335L | This work |
| *E. coli* BL21 pYT/his-*glnA3* | pYT9 derivative for overexpression of His-*glnA3*<sub>Sc</sub> | (16) |
| *M. tuberculosis* H37Rv | wild type | (54) |
| *M. tuberculosis* H37Rv $\Delta glnA3$ | *M. tuberculosis* derivative with deleted *glnA3* gene | This work |
| **Plasmids** |  |  |
| pRM4 | pSET152 *ermEp* * with artificial RBS | (52) |
| pRM4*glnA3* | pRM4-derivative with *glnA3*<sub>Sc</sub> (*S. coelicolor*) | (48) |
| pRM4*glnA3*<sub>Mt</sub> | pRM4-derivative with *glnA3*<sub>Mt</sub> (*M. tuberculosis*) | This work |
| pET30 Ek/LIC | ligation independent cloning (LIC), T7 Promoter, T7 transcription start, phage f1 origin of replication, detachable N-terminal His-tag and S-tag, C-terminal His-tag, T7 terminator, *lacI* coding sequence, pBR322 *ori* | Novagen |
| pET30*glnA3*<sub>Mt</sub> | pET30-derivative with *glnA3*<sub>Mt</sub> (*M. tuberculosis*) | This work |
| pK18 | pUC-derivative, *lacZ'*α-complementation system, Km<sup>R</sup> | (53) |

**Table 4.**
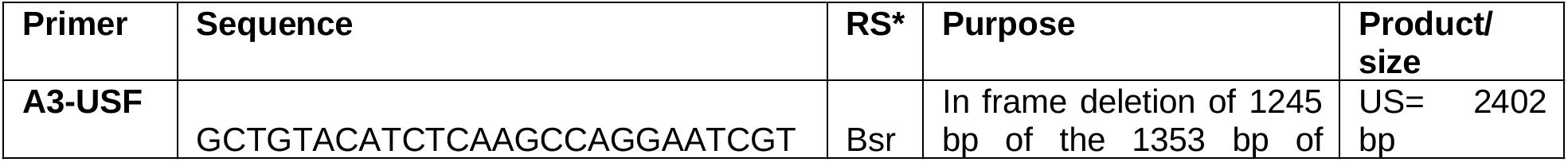
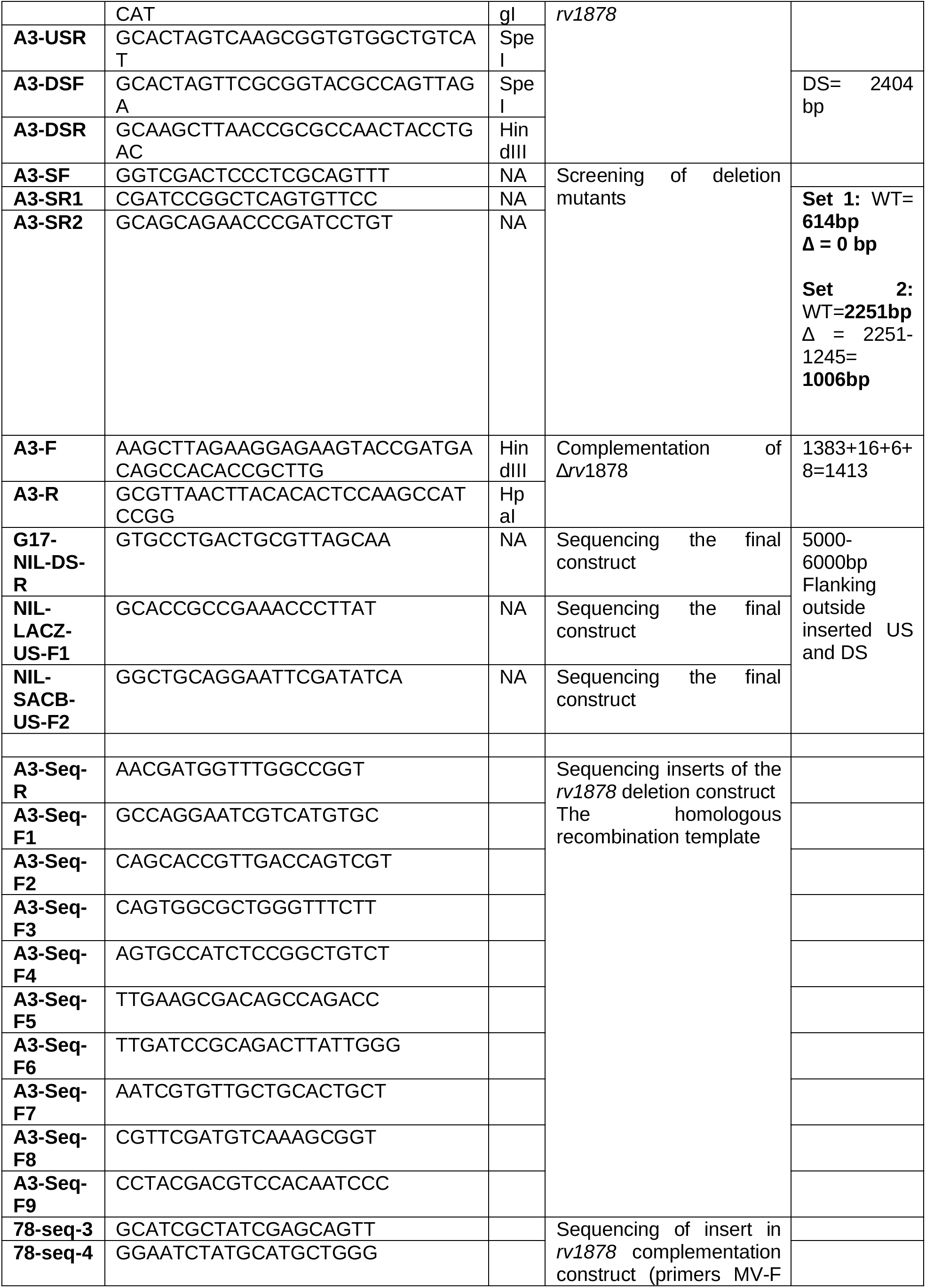
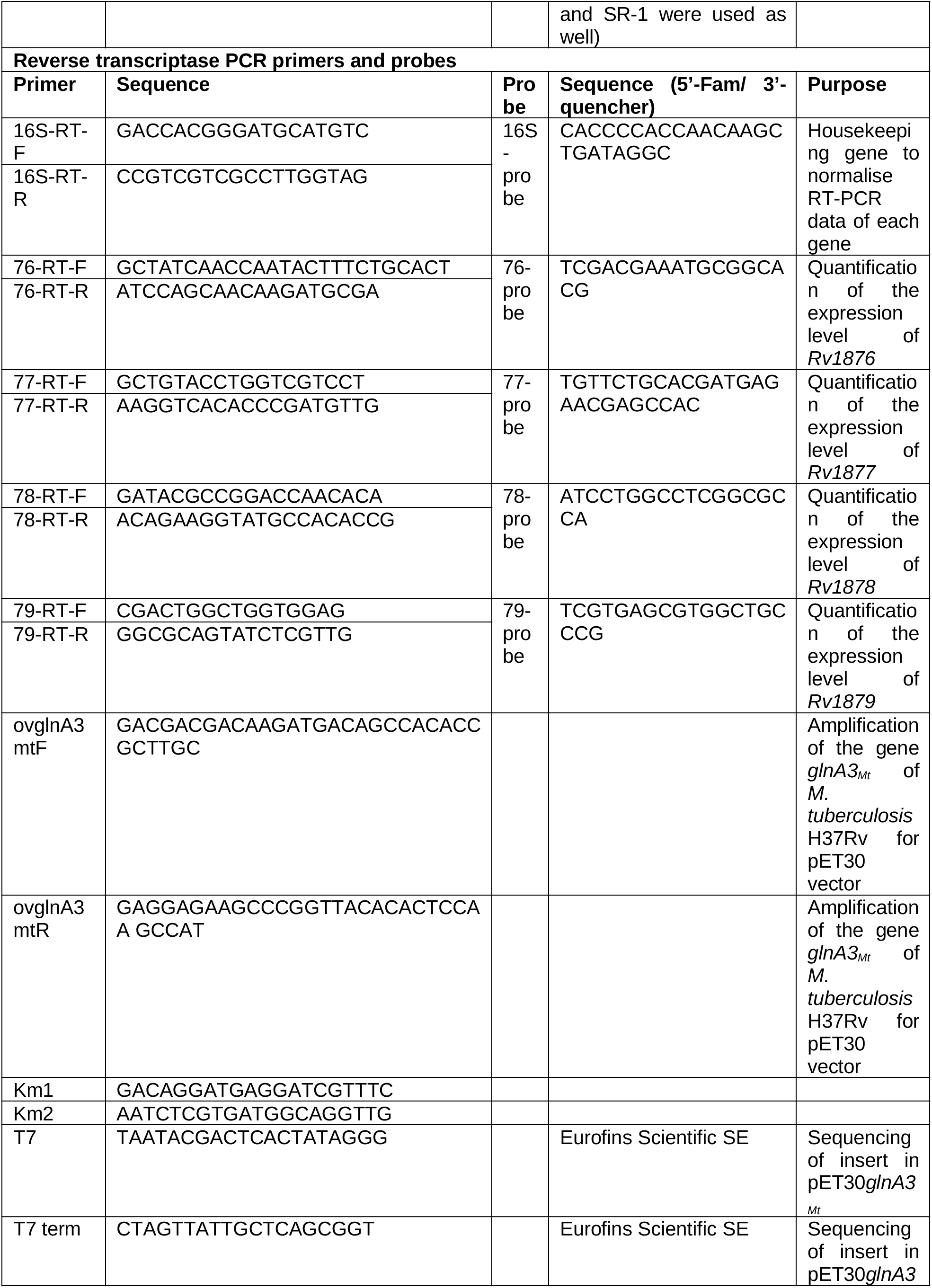
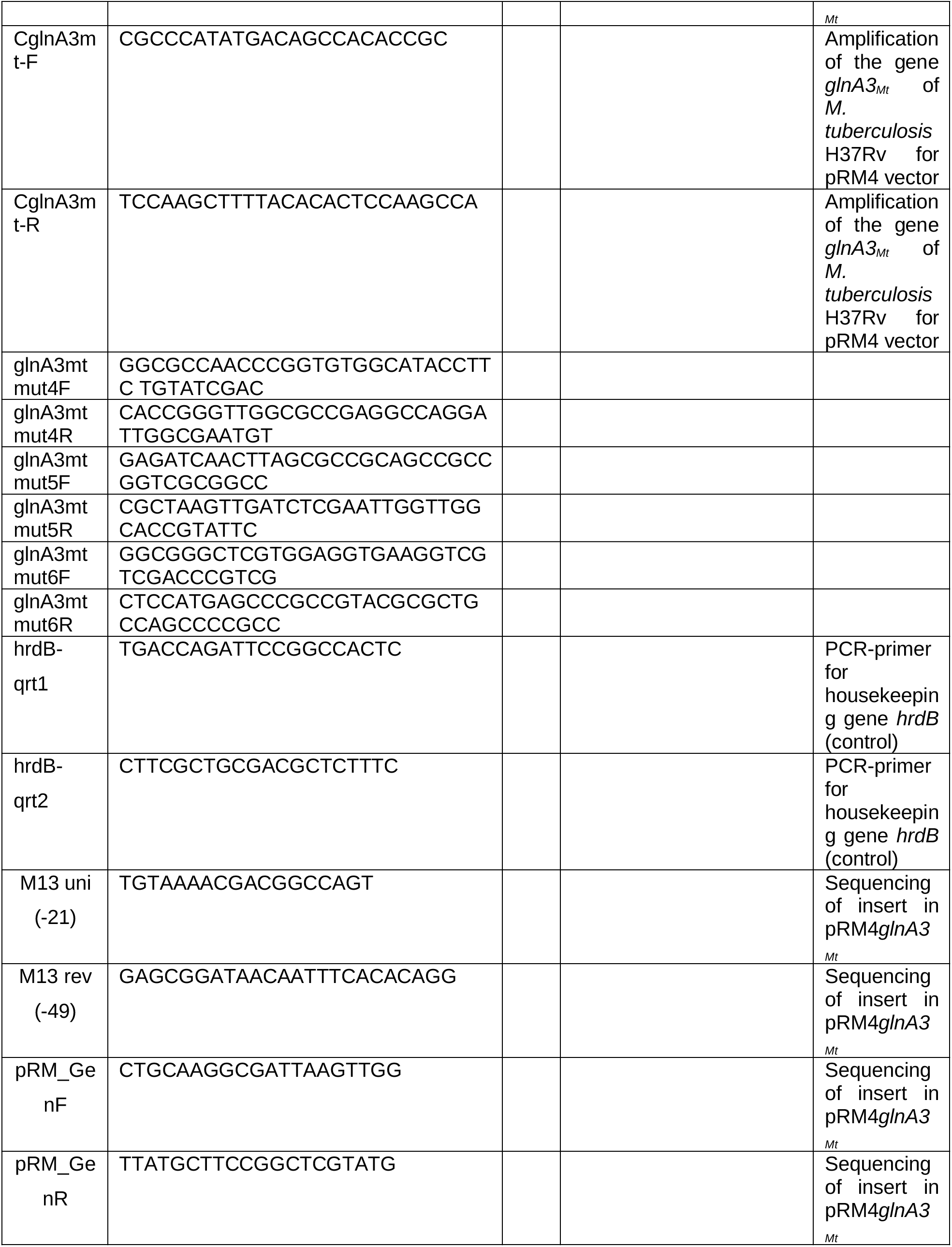
Primers used in the study.

### Complementation of the *S. coelicolor ΔglnA3* mutant

For complementation of the *S. coelicolor glnA3_Sc_* mutant, the *glnA3_Mt_* gene from *M. tuberculosis* was amplified by PCR using *CglnA3mtF* and *CglnA3mtR* primers and cloned into the multiple cloning site of pRM4 plasmid between *Nde*I and *Hin*dIII restriction sites downstream of the constitutively expressed erythromycin promoter *ermE*p* located on the plasmid. The kanamycin resistance cassette was amplified by PCR using the pK18 plasmid as a template and primers *aphIIupperEcoR*I as well as *aphIIlowerHind*III, and introduced to the multiple cloning site of the recombinant plasmid pRM4-*glnA3MtC* between *Eco*RI and *Hin*dIII restriction sites. The correct construct was confirmed by colony PCR as well as by sequencing and introduced into the *S. coelicolor glnA3_Sc_* mutant by biparental conjugation using *E. coli* S-17. Clones were selected on resistant phenotype against kanamycin and apramycin. The correct integration of the pRM4-*glnA3Mt* was confirmed by PCR and sequencing.

### Cloning, expression, and purification of His-Strep-GlnA3*_Mt_*

The GlnA3*_Mt_* encoding gene *Rv1878* was amplified using genomic DNA of *M. tuberculosis* as template. It was inserted into the expression vector pET-30 Ek/LIC (Novagen) under the control of the IPTG inducible T7 promoter using the pET-30 Ek/LIC Cloning Kit (Novagen). His/Strep-GlnA3_Mt_ was synthesized in *E. coli* BL21 (DE3). Initially cells were incubated over night at 37°C in LB medium, afterwards transferred in fresh LB medium and incubated for at 25°C until the culture reached an optical density of 0.5 at 600 nm. Subsequently, the culture was induced with 1 mM IPTG and incubated at 25°C overnight. His/Strep-GlnA3_Mt_ was purified by nickel ion affinity chromatography essentially as directed by the resin manufacturer (GE-Healthcare). Purified His/Strep-GlnA3_Mt_ was dialyzed against 20 mM Tris, 100 mM NaCl (pH 8.0) or used for further purification steps. His/Strep-Tag was cleaved using enterokinase according to the protocol of the manufacturer (NEB) and GlnA3*_Mt_*was immediately purified from the digestion mix by size-exclusion chromatography as directed by the resin manufacturer (GE-Healthcare).

### HPLC/ESI-MS detection of a glutamylated product

For the detection of the glutamylated product of the GlnA3*_Mt_*catalyzed reaction, an HPLC/ESI-MS procedure has been applied. Standard reactions contained: 20 mM HEPES (pH 7.2), 10 mM ATP, 150 mM glutamate sodium monohydrate, 150 mM putrescine dihydrochloride, cadaverine dihydrochloride, spermidine trihydrochloride, or spermine tetrahydrochloride, 20 mM MgCl_2_x6H_2_O, were mixed with 10 µg of the purified His/Strep-GlnA3*_Mt_* or GlnA3*_Mt_* (or without GlnA3 as a control) and incubated at 30°C for 5 min. The reaction mixture was incubated at 100°C for 5 min in order to stop the reaction. HPLC/ESI-MS analysis was performed on an Agilent 1200 HPLC series using a Reprosil 120 C_18_ AQ column, 5 µm, 200 mm by 2 mm fitted with a precolumn 10 mm by 2 mm (Dr. Maisch GmbH, Ammerbuch, Germany) coupled to an Agilent LC/MSD Ultra Trap System XCT 6330 (Agilent, Waldbronn, Germany). For analysis were used: 0.1% formic acid as solvent A and acetonitrile with 0.06% formic acid as solvent B at a flow rate of 0.4 ml min^-1^. The gradient was as follows: t_0_ = t_5_ = 0% B, t_20_ = 40% B (time in minutes). Injection volume was 2.5 µl, column temperature was 40°C. ESI ionization was done in positive mode with a capillary voltage of 3.5 kV and a drying gas temperature of 350°C.

### Modified GS activity assay

The enzymatic activity of GlnA3_Mt_ variants was tested in a modified GS activity assay(24). Solutions A, B, C and F and the reaction mix were prepared containing enough protein for the release of 35-50 mM Pi produced in 5 min. After the adjustment of the pH, 95 μl were loaded into PCR-strips for each reaction. Solution D or Solution E for undetermined kinetic parameters of ATP was prepared. Afterwards, the reaction was initiated by adding 5 μl substrate to the reaction mix. Additionally, blanks with H_2_O_deion_ and a phosphate standard ranging from 0 to 20 mM were included. The reaction mix was incubated at 30°C for 5 min in a thermocycler. The wells of a 96-well plate were loaded with 150 μl of solution D (or C) for each reaction. Afterwards, 50 μl of the reaction mix was transferred to the previously prepared solution D (or C) in the 96-well plate. Solutions were mixed and incubated for 5 min at RT. At low pH the enzymatic reaction was terminated, while 150 μl of solution F were added to stop color development. The final reaction was incubated for 15 min at RT. The absorbance was measured at 655 nm using a Microplate reader. Raw absorbance readings were put into Excel (Microsoft).

### *In silico* protein modeling and docking studies

To build the *in silico* models for the different GlnA-like enzymes an existing template from the Protein Data Bank (PDB) was selected. The initial template search was done using the template search function of SWISS-MODEL (25) by input of the amino acid sequence in FASTA format, plain text or UniProtKB accession code. SWISS-MODEL was then performing search for evolutionary related protein structures against the SWISS-MODEL template library SMTL (60) and using database search methods BLAST, and HHblits (61, 62). The resulting template structures were ranked and further evaluated by SWISS-MODEL using the estimated Global Model Quality Estimate (GMQE) (60) and the Quaternary Structure Quality Estimate (QSQE) (63). Top-ranked templates and alignments were compared to verify whether they represent alternative conformational states or cover different regions of the target protein. In such case, multiple templates were selected automatically and different models were built accordingly (25). Out of the resulting list of possible templates different templates were chosen based on the GMQE, QSQE, identity and oligo state. Also, value was given to select templates out of different bacterial phyla to get quality control with diversity. Using the selected templates as a base SWISS-MODEL built a 3D protein model estimating the real 3D structure of the protein. Therefore, SWISS-MODEL started with the conserved atom coordinates defined by the target-template alignment and then coordinated residues corresponding to insertions/deletions in the alignment that were generated by loop modelling and a full-atom protein model was obtained by constructing the non-conserved amino acid side chains (25). SWISS-MODEL used the ProMod3 modelling engine and the OpenStructure computational structural biology framework (64). The evaluation of the build 3D protein models was done using the QMEAN scoring function, the MolProbity score and a Ramachandran plot of the model. The QMEAN score provided an estimate of the ‘degree of nativeness’ of the structural features observed in a model and described the likelihood that a given model was of comparable quality compared to experimental structures (31). The MolProbility score relied heavily on the power and sensitivity provided by optimized hydrogen placement and all-atom contact analysis, complemented by updated versions of covalent-geometry and torsion-angle criteria (33). The Ramachandran plot plotted the torsion angles of the different amino acids against each other to verify the correct folding of the *in silico* model (32). Based on the highest QMEAN score and the Ramachandran plot the best *in silico* 3D protein models were chosen. Molecule structures were obtained from the PubChem database. Molecular docking was performed using UCSF Chimera software.

### Site directed mutagenesis

To select possible key amino acids the *in silico* 3D structures of GlnA3*_Mt_* were compared with the template structures they originate from. The main used 3D model of the GlnA3*_Mt_* enzyme was based on the GlnA1 (PDB code 1HTO) structure of *M. tuberculosis* (26). Therefore, the 3D structure model of GlnA3*_Mt_*and the template structure were opened with SWISS-PDB viewer (65), PyMol (PyMOL, The PyMOL Molecular Graphics System, Version 2.0 Schrödinger, LLC.) or UCSF Chimera. Through literature research key amino acids, of the template, could be identified and then marked in the 3D template structure. The amino acids that overlap directly with the key amino acids of the template were marked. The used site-directed mutagenesis approach was based on the protocol described by Zheng et al. (66). It was using a PCR approach with specifically designed primers to induce into plasmid DNA. The primers contained the wanted mutation with was introduced in the plasmid DNA through the PCR amplification. The GlnA3*_Mt_*\* mutated variants were generated using the pET-30 Ek/LIC plasmid with the cloned *glnA3* gene as template and were expressed in *E. coli* strain BL21 (DE3).

### RNA sample preparation

A volume of 10 ml of early to mid-logarithmic phase cultures was either treated with spermine or the DMSO control, and harvested 3 hours later using the RNA Pro Blue kit (MP Bio), and the resuspended cells were homogenised using the Fast Prep Homogenizer (30 seconds, 6 m/s, four times, 5 min intermittent). The cell lysate was filtered twice using PTFE syringe filters (13 mm diameter, 0.2 µM pores size) and taken out of the BSL3 laboratory for further purifications. The first purification was performed using the Direct-zol RNA Miniprep Plus (R2070, 100 µg binding capacity) including an in-column DNA digestion step, according to the manufacturer’s instructions. The purified samples were quantified using a spectrophotometer, and diluted if the concentration exceeded 200 ng/µl. Then they were further digested using the Turbo DNA-free kit (Thermofischer) according to manufacturer’s instructions, however in two consecutive rounds, to ensure complete DNA digestion. The digested samples were further purified and concentrated using the RNA clean and concentrator kit-25 (R1017, 50 µg binding capacity), including another in-column DNA digestion step, according to manufacturer’s instruction. The resulting samples were used for Next Generation RNA sequencing performed by Eurofins Genomics GmbH.

### RNA sequencing

Bulk RNA sequencing of the resulting samples as well as expression quantification was performed by Eurofins Genomics GmbH. RNA quality was measured using the Agilent 2100 Bioanalyzer and all but one sample had RIN value larger 8. Sequencing was performed on the Illumina Novaseq 6000 using the NEBNext(R) Ultra II Directional RNA Library Prep Kit, generating between 13.5 and 17.3 million 150-bp paired-end read pairs. Reads were mapped to the genome sequence of Mycobacterium tuberculosis H37Rv (NC_000962.3; obtained from https://www.ncbi.nlm.nih.gov/nuccore/NC_000962.3?report=fasta) using bwa-mem version 0.7.12-r1039 (67). With the exception of one sample with 89% of reads mapping to H37Rv, all samples had a mapping rate larger than 97%. Reads were subsequently counted for 3,906 *Mycobacterium tuberculosis* H37Rv genes according to RefSeq annotation (obtained via https://www.ncbi.nlm.nih.gov/Taxonomy/Browser/wwwtax.cgi?id=83332) using tool featureCounts (68). Raw overall read counts per sample are between 4.4 and 8.8 million (counting each read pair once and omitting multi-mapping and low-quality reads). Trimmed mean of M-values (TMM) normalization (69) of raw counts was performed using the edgeR package version 3.16.5 (70). In a multidimensional scaling plot, the first dimension separated spermidine-treated from untreated samples. Normalized counts were used for computation of fold changes and computation of median fold changes, which were log2 transformed whenever specifically noted.

### RT-PCR

In order to perform the reverse-transcriptase quantitative PCR, (RT-PCR), RNA samples were converted to cDNA using the Maxima First Strand cDNA KIT (Thermofischer). During method optimization, controls where all reagent were added except the reverse transcriptase were included (non-reverse transcriptase control), in order to run it along with the converted samples, to ensure that genomic DNA was completely removed or negligible. The reverse transcribed samples were run in a 10 µl reaction on a LightCycler 480, using the LightCycler 480 master mix. Quantification was made by the integrated software of the LightCycler, according to a probe-based assay (labelled at the 5’-end with FAM and at the 3’-end with a quencher), using *M. tuberculosis* genomic DNA to generate a standard curve (10-1000 pg/µl). The probe and primers were designed by TIB MOLBIOL Syntheselabor GmbH. Various set of primers were designed, and these were tested, to obtain the optimal set, that was used for the quantification (Table 3).

## 4. DISCUSSION

During evolution, many intracellular pathogens (including protozoan parasites) have developed strategies to access selected nutrients from the host for their growth and survival. *M. tuberculosis* is an example of an intracellular pathogen, which is very well adapted to survive within the hostile macrophage environment. The long term host and pathogen persistence is the consequence of a subtle equilibrium between the nutritive needs of host and pathogens. On one hand, the presence of high polyamine levels in the M2 intracellular environment of macrophages may lead to growth inhibition or cell death of *M. tuberculosis* (71). Already 70 years ago it was reported that polyamines like spermine show a tuberculostatic activity (72). On the other hand, polyamines may provide a nutrient source that can be exploited by *M. tuberculosis*.

Since free spermine is toxic to cells, it has to be immediately excreted or metabolized further and neutralized by a modification such as acetylation (73) or gamma glutamylation (16, 18). The modified spermine is no longer toxic for bacteria due to loss of its positive charge (74). Glutamylated spermine may be subsequently utilized as N/C-source or excreted from cells.

In order to investigate how *M. tuberculosis* can survive under excess polyamine we took advantage of the knowledge on polyamine metabolism in *S. coelicolor,* which is a model actinobacterium that shares essential features in nitrogen metabolism with *M. tuberculosis*. The gamma-glutamylpolyamine synthetase GlnA3*_Sc_* is the primary enzyme of the polyamine utilization pathway in *S. coelicolor* (16) catalyzing the first step of polyamine utilization, the detoxification of polyamine by glutamylation. Thus, we hypothesized that a similar polyamine utilization pathway might enable pathogenic actinobacteria, such as *M. tuberculosis*, to colonize host cells and evade the host’s immune response by conferring resistance against toxic polyamines and ensuring putative polyamine utilization necessary for long-term persistence. Transcription of *glnA3_Mt_*, the mycobacterial orthologue of *glnA3_Sc_*, was reported in broth culture (75) and the respective protein was detected in a guinea pig model of tuberculosis *in vivo*, during the chronic stage of this disease (76, 77). *glnA3_Mt_* was reported to be poorly expressed in *M. tuberculosis* under laboratory conditions (19). Its presence in the cell is not essential for bacterial homeostasis, however its role was not evaluated in the presence of high polyamines concentrations reflecting disease relevant conditions.

In the current study we observed that *M. tuberculosis* growth can be inhibited by spermine. Using a GS-based *in vitro* enzymatic activity assay we show that GlnA3*_Mt_* (Rv1878) acts as a gamma-glutamylspermine synthetase and generates a glutamylated spermine. In an *in vitro* phosphate release assay we demonstrated that purified recombinant GlnA3*_Mt_* prefers spermine as a substrate, over putrescine, cadaverine, spermidine or other monoamines and amino acids. These results lead to the conclusion that GlnA3*_Mt_* may play a specific role in the detoxification of the polyamine spermine. We therefore speculated that GlnA3*_Mt_* may be essential for the survival of *M. tuberculosis* during spermine stress. However, the deletion of the *glnA3* gene in *M. tuberculosis* did not result in immediate death of the bacteria or a reduced growth rate of the strain in the presence of high spermine concentrations. Thus, at the moment, the specific functional role of GlnA3*_Mt_* in *M. tuberculosis* remains unclear. The assumption that GlnA3 may play a detoxifying role only at rather low spermine levels could be excluded since a similar growth rate of WT and *glnA3-*deficient *M. tuberculosis* was observed at all spermine concentrations tested.

A further finding that GlnA3_Mt_ is probably not crucial for survival under polyamine stress was obtained from transcriptional analyses: The *gln*A3*_Mt_*gene is part of a locus consisting of four genes, comprising (*rv1876*) – encoding bacterioferritin A (*bfrA*) (77), *rv1877* – encoding a multidrug efflux pump (40), *glnA3* (*rv1878*) - gamma glutamyspermine synthetase, and *rv1879* – encoding a putative amidohydrolase. Both RNAseq analyses and subsequent confirmatory RT-PCR showed that the gene cluster *rv1876-rv1877-rv1878-rv1879* was not upregulated in the presence of spermine, suggesting that GlnA3_Mt_ does not participate in a spermine-induced transcriptional response in this bacterium.

In contrast, RNAseq revealed three alternative genes to be significantly upregulated in response to spermine: *rv3065*, *rv3066*, and *rv3067*. Interestingly, *rv3065* encodes an efflux pump suggesting that active excretion of polyamines is the bacterium’s primary defence against this class of molecules. Rv3065 was the first mycobacterial protein identified and described in the small membrane protein family (SMR) in *M. tuberculosis* (78). This represents a group of efflux pumps that contain four transmembrane domains and confer resistance to aromatic dyes, derivatives of tetraphenylphosphonium (TPP) and quaternary amines (79). Rv3065 has independently been shown to mediate the efflux of different chemical compounds and antibiotics belonging to the pyrrole and pyrazolone chemical classes (80). Over-expression of *rv3065* in *M. tuberculosis (*81) and in *M. smegmatis* (78) increased resistance to to tetraphenylphosphonium (TPP), erythromycin, ethidium bromide, acriflavine, safranin O, and pyronin Y.

Of note, joint transcription of *rv3065*-*rv3067* has previously been observed in *M. tuberculosis* treated with thioridazine (80). Thus, it is possible that spermine may also be a substrate for Rv3065. We have also seen that *rv1877*, encoding for another efflux pump was up-regulated during spermine stress in 7H9 medium, thereby pointing also to a particular role of efflux activities. Such a resistance mechanism would make the detoxification of spermine superfluous and could explain the survival of the *glnA3Mt* mutant in the presence of spermine

### Conclusion

In this study, the enzymatic function of GlnA3*_Mt_* and the anti-mycobacterial activity of spermine were demonstrated. However, further studies are needed to decipher the spermine detoxification mechanisms in *M. tuberculosis*. In particular, an interplay of GlnA3 and the second gamma-glutamylpolyamine synthetase GlnA2 as well as potential polyamine transporter Rv3065 seems to compose a complex system to escape spermine toxicity by *M. tuberculosis*.

## ACKNOWLEDGEMENTS

We acknowledge support by the Federal Ministry of Education and Science (Bundesministerium für Bildung und Forschung - BMBF) and cooperation partners from the GPS-TBT and GSS-TUBTAR projects as well as the DZIF projects within TTU-TB “New drugs and regimens” and the TTU-NAB “Novel Antibiotics”. WW acknowledges infrastructural funding from the Cluster of Excellence EXC2124 (Controlling Microbes to Fight Infection, project ID 390838134) from the DFG. We gratefully acknowledge Carolin Golin (Borstel) and Alena Strüder (Tübingen) for expert technical assistance and Dr. Gareth Prosser (Borstel) for helpful suggestions and critical reading of the manuscript. We acknowledge support by Open Access Publishing Fund of University of Tübingen.

## 5. FUNDING

This research was funded by the Federal Ministry of Education and Science BMBF (Fördermaßnahme “Targetvalidierung für die pharmazeutische Wirkstoffentwicklung”), project GPS-TBT (FKZ: 16GW0183K, 16GW0184) as well as project GSS-TUBTAR (FKZ: 16GW0253K, 16GW0254).

## 6. CONFLICT OF INTERESTS

All authors declare no conflict of interest.

## 7. AUTOR CONTRIBUTIONS

AM, WW, NR, DH, CSE, FH, SK designed the study and developed the methodology. SK and MO cloned, overexpressed and purified the GlnA3*_Mt_* protein. SK and MB cloned, overexpressed and purified the His-Strep-GlnA3*_Mt_*\* protein variants. SK and MO constructed and analyzed the *ΔglnA3_Sc_*pRM4*glnA3_Mt_* complementation strain. SK analyzed DNA/protein sequences, protein structures, molecular docking data, HPLC, MS/MS and assay data. CSE generated the *M. tuberculosis glnA3* mutant, prepared *M. tuberculosis* samples, performed survival assays, carried out RT-PCR analysis and prepared and managed RNAseq samples and results. IW investigated RNA-Seq data analyses and computed and visualized expression changes. SK, CSE, NR wrote the manuscript. SK and MB performed homology modeling of proteins. SK, MO and MB performed phenotypical analysis of all mutants and parental strain *S. coelicolor* M145 on Evans medium with different nitrogen sources. CM cloned, overexpressed and purified SUMO-GlnA3*_Mt_*\* protein variant. SK, MB, and CM performed GlnA3*_Mt_ in vitro* assay. AK performed the HPLC and HPLC/MS analyses. WW, NR, DH, CSE, FH, HBO provided helpful feedback on an early draft of the paper and assisted with data analysis. SK, MO, MB, NR and CM prepared all figures for manuscript. SK, CM, AM, AK, WW, NR, DH, FH, and IW contributed to the editing of the manuscript and resolved final approval of the version to be published.

## Supplementary figures

**Suppl. Fig. 1.**
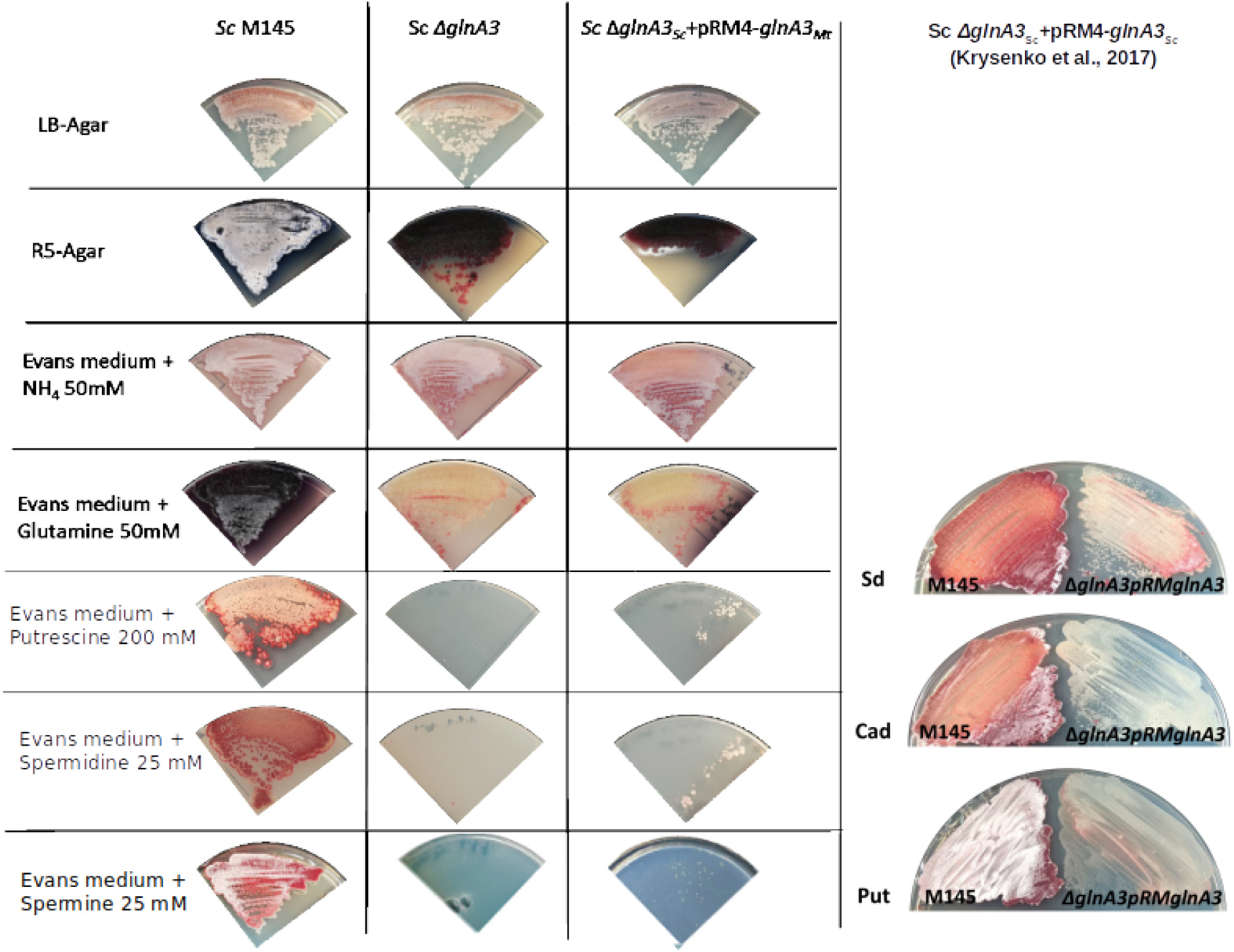
Physiological role of the *glnA3_Mt_* gene product in the *S. coelicolor glnA3* mutant grown in the presence of polyamines and other nitrogen sources. Phenotypic comparison of the parental strain *S. coelicolor* M145, the *glnA3* mutant and the *ΔglnA3Sc*pRM4*glnA3_Mt_* complementation strain grown on rich LB-agar, R5-agar or defined Evans medium supplemented with glutamine (50 mM), NH_4_Cl – ammonium chloride (50 mM) and polyamines putrescine (200 mM), spermidine or spermine (25 mM) as sole nitrogen source. Each panel represents observations on a single agar plate. Complementation of the *S. coelicolor glnA3* mutant with *glnA3_Mt_* resulted in restored growth of the strain in the presence of high putrescine, spermidine and spermine concentrations.

**Suppl. Fig. 2.**
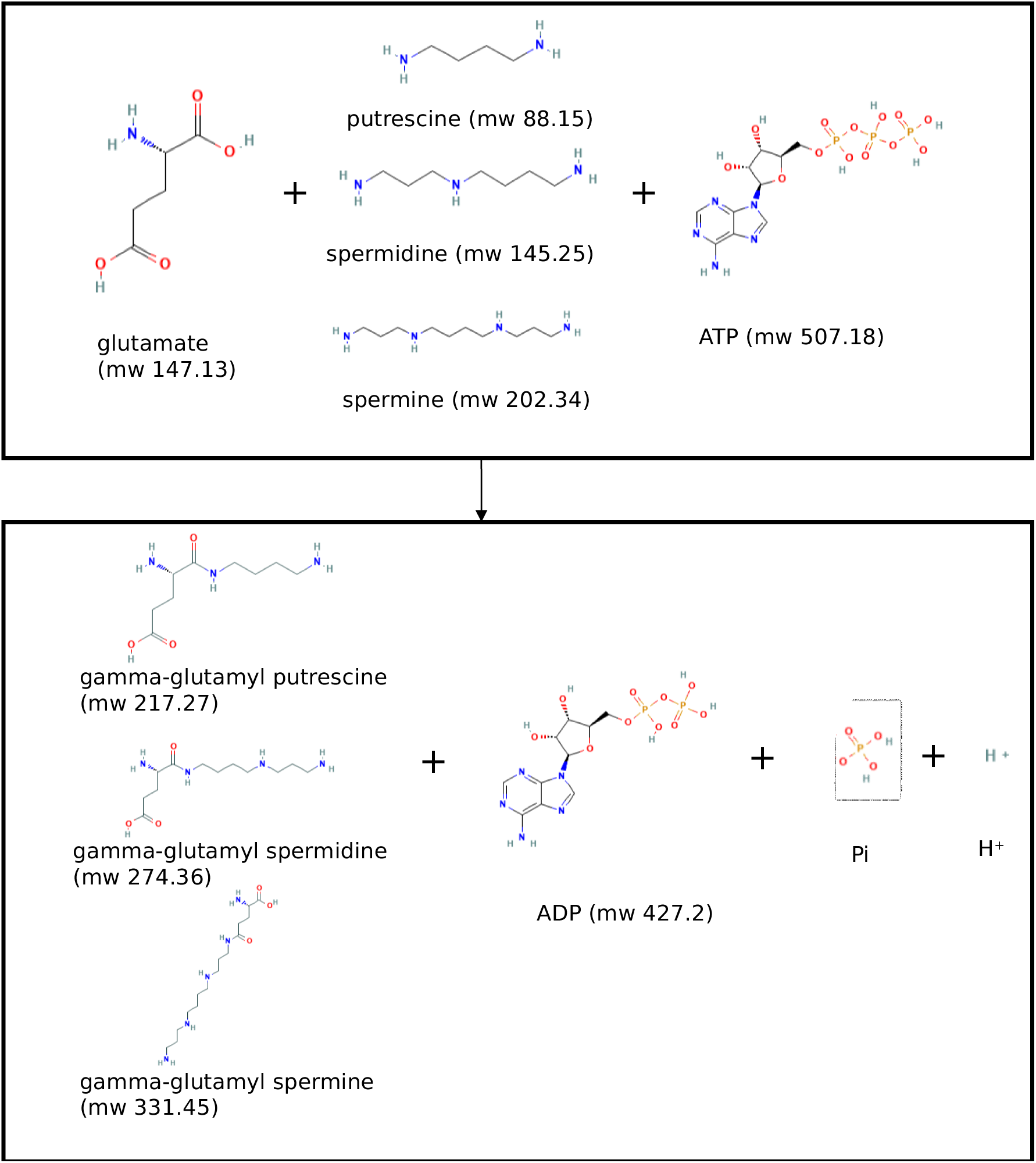
Reaction catalyzed by gamma-glutamylpolyamine synthetase with indicated substrates, products and molecular weight of each molecule.

**Suppl. Fig. 3.**
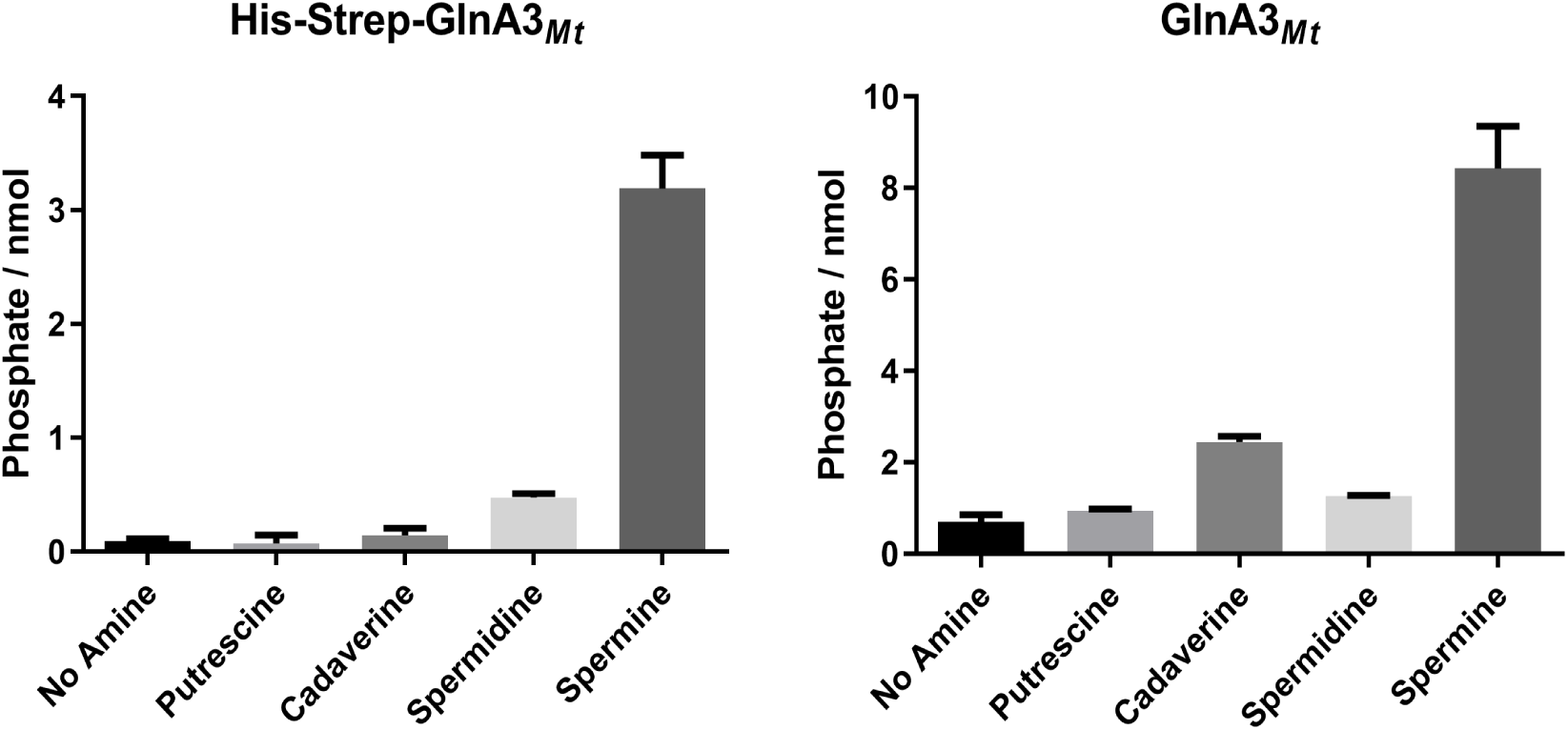
Specific activity of the native GlnA3*_Mt_* with different polyamines. All substrates are at a concentration of 50 mM. The mean value of minimum n=3 biological replicates with n=3 technical replicates each with standard error is shown.

**Suppl. Fig. 4.**
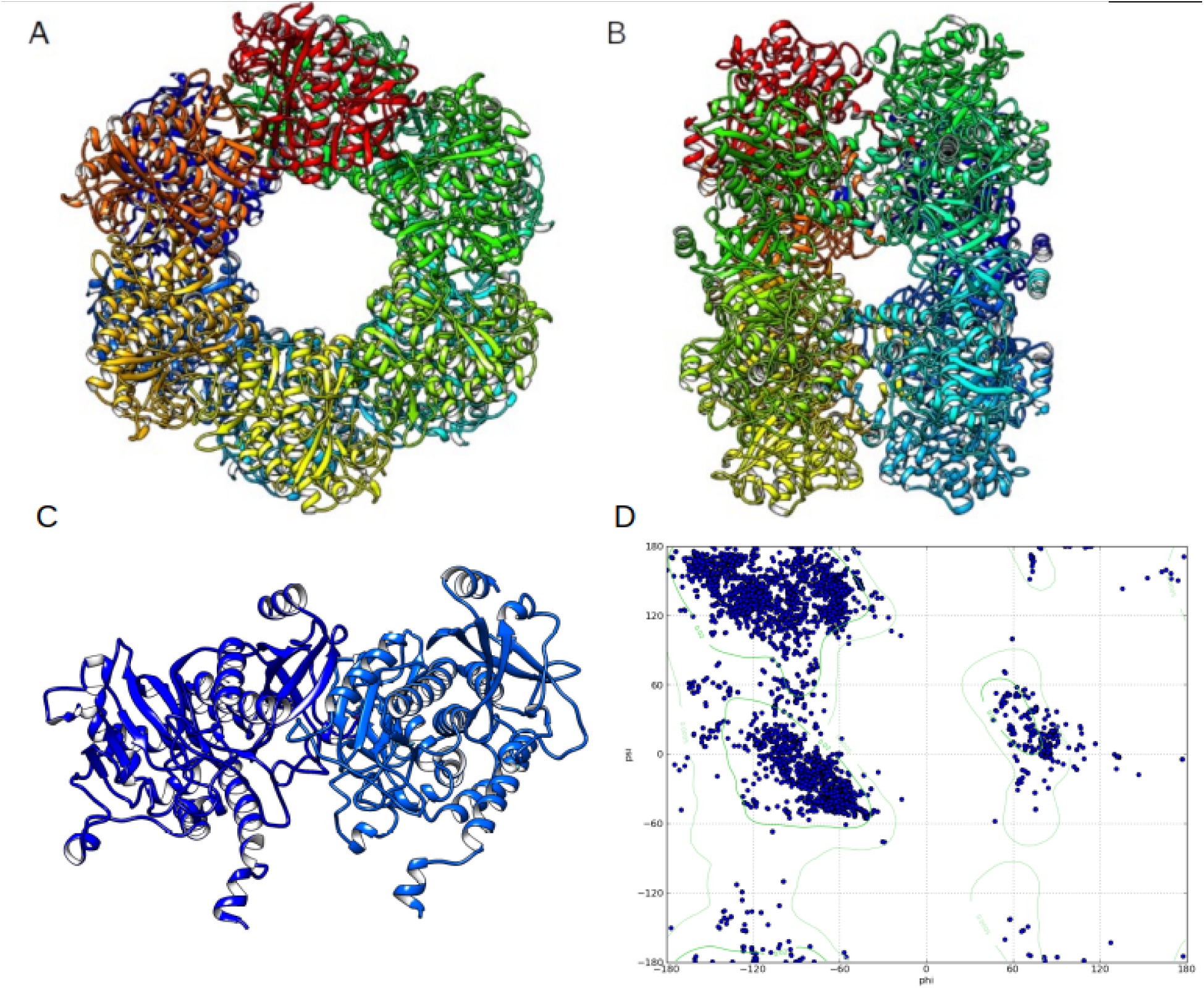
Structural alignment of the 3D model structure of GlnA3*_Mt_*, based on and superposed with the GlnA*_Mt_* template (PDB code 1BVC (27)). A) each of the twelve subunits is marked in a different color (color scheme “chain”), B) side-view on two 6-unit containing rings, C) a representation of the active site and substrate binding pocket, D) Ramachandran plot demonstrating the good quality of the generated GlnA3*_Mt_* model, based on visualization of energetically allowed regions for backbone dihedral angles ψ against φ of amino acid residues in the structure.

**Suppl. Fig. 5.**
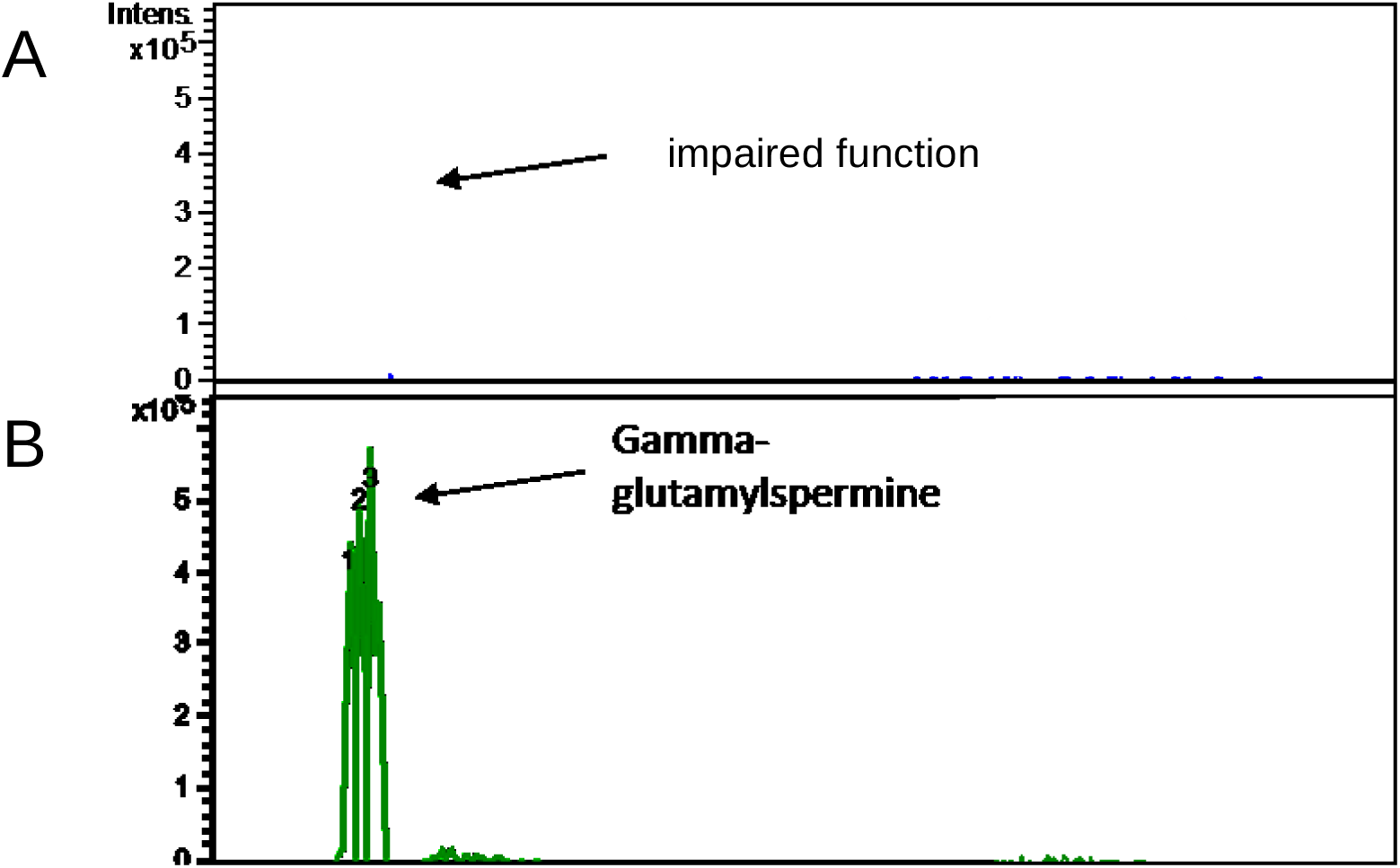
HPLC/ESI-MS analysis of His-Strep-GlnA3*_Mt_*SER199 and His-Strep-GlnA3*_Mt_*. Two samples were analyzed in MS negative mode: reaction mixtures with addition of His-Strep-GlnA3*_Mt_*SER199 (A) and with addition of His-Strep-GlnA3*_Mt_* (B). Extracted ion chromatograms for the His-Strep-GlnA3*_Mt_* reaction product corresponding to gamma-glutamylspermine with charge to mass ratio of *m/z* 331 was shown (B), and no product in the sample with GlnA3*_Mt_*SER199 was detected (A).

**Suppl. Fig. 6.**
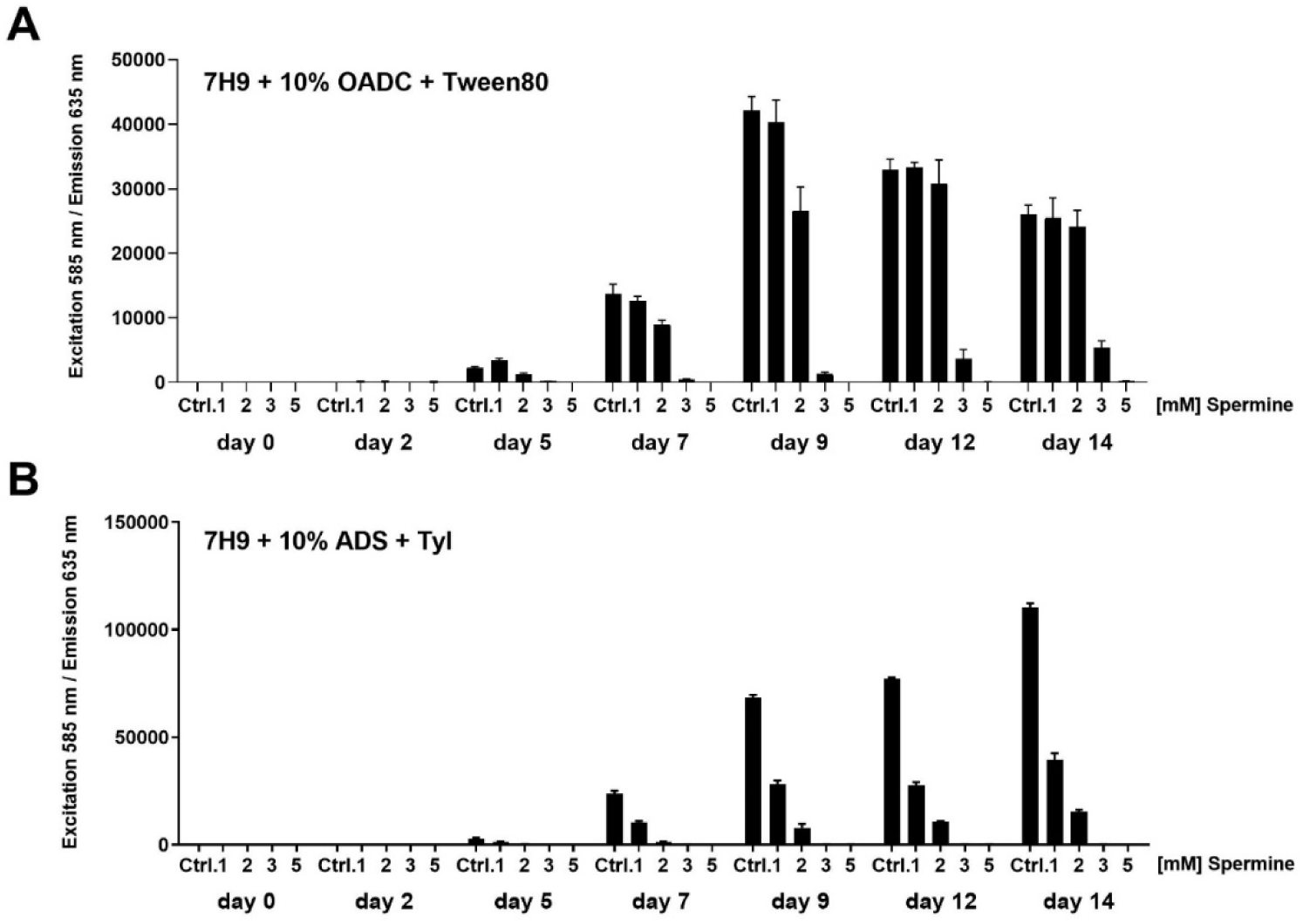
Growth of *M. tuberculosis* in the presence of spermine. *m*Cherry10-expressing Mtb H37Rv bacteria (55) were incubated for a period of 14 days in the absence or presence of spermine in (A) 7H9 medium + OADC + Tween80 or (B) in 7H9 medium + ADS + Tyloxapol and analyzed as described (55); Ctrl: DMSO; Spm: spermine. Concentrations tested were from 1mM to 5 mM.

**Suppl. Fig. 7.**
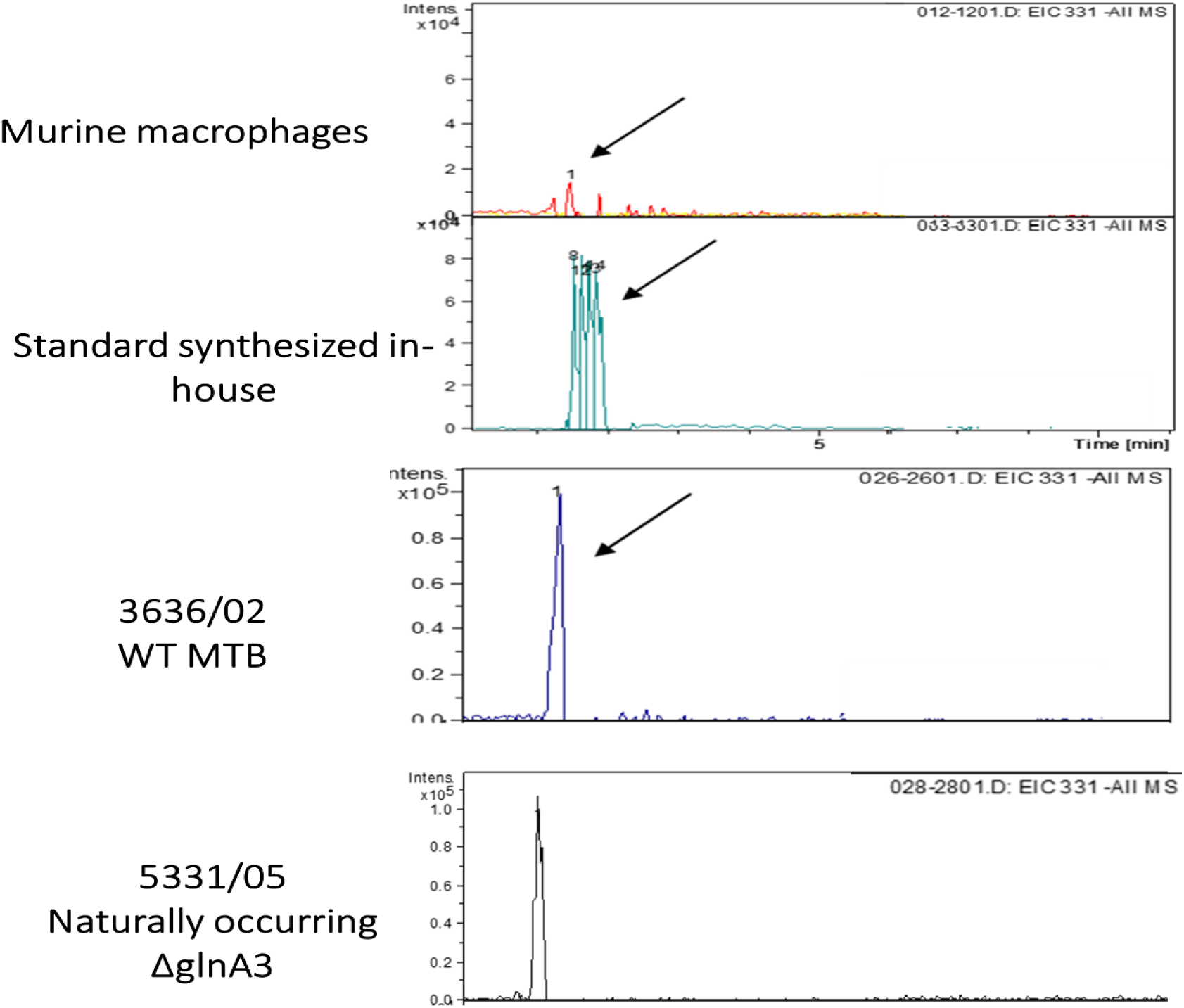
HPLC/ESI-MS detection of gamma-glutamylspermine in the MTB Beijing strain (WT variant) and in the naturally occurring *glnA3* Beijing mutant. Samples were analyzed in MS negative mode.

**Suppl. Fig. 8.**
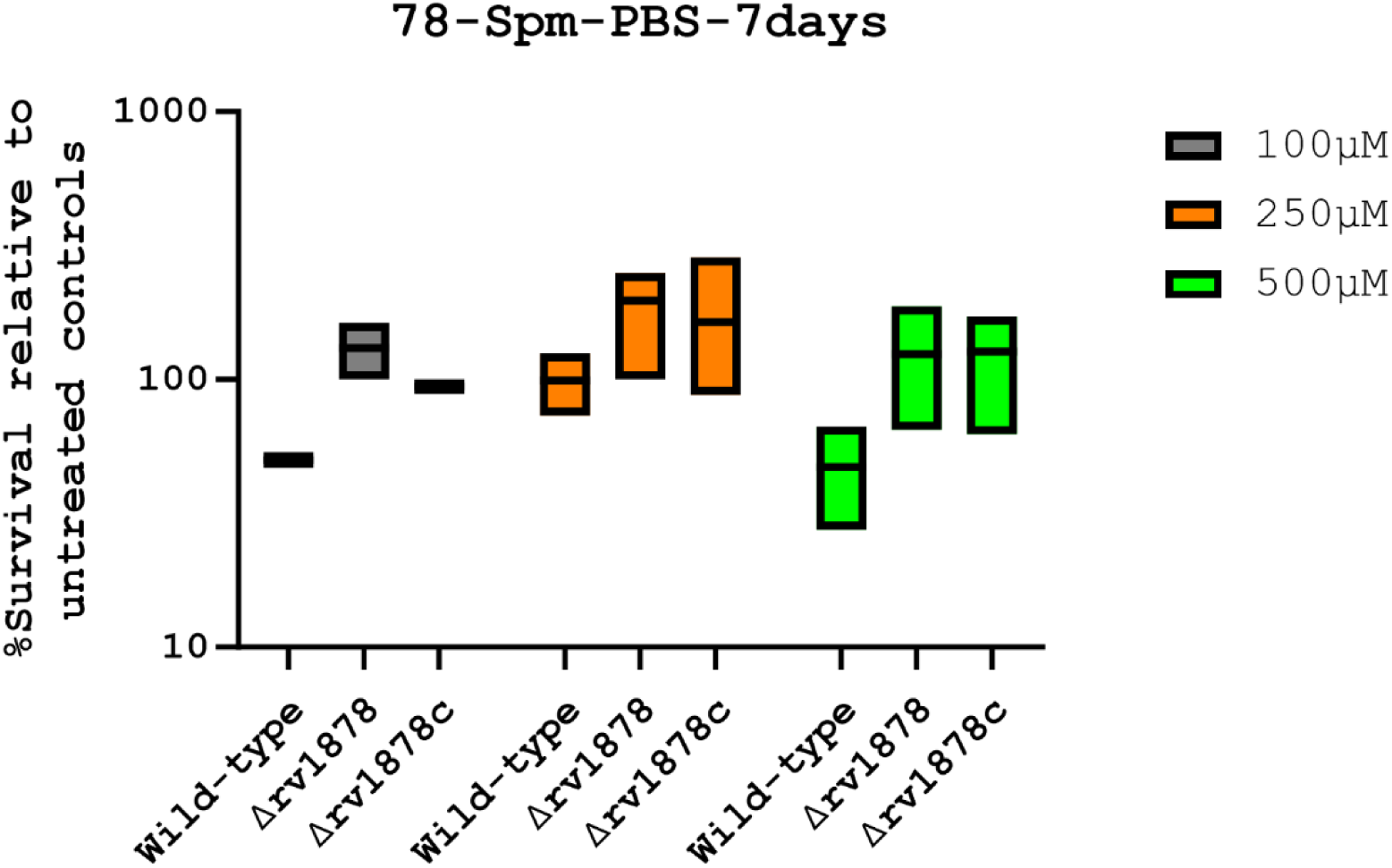
Spermine enhances the survival of the Δ*glnA3* mutant as a sole C/N source.Logarithmic phase cultures (in Sauton’s media), were washed several times in PBS, diluted to approximately 10^5^ CFUs/ml, and incubated in the presence (treated) or absence (untreated) of various concentrations of spermine over 7 days. Survival was determined by dividing the number of colonies obtained (after plating strains from various conditions) from the treated conditions to the untreated condition.

## Notes

### Competing Interest Statement

The authors have declared no competing interest.

