## Supplemental 1 for "GlnA3*_Mt_* is able to glutamylate spermine but it is not essential for the detoxification of spermine in *Mycobacterium tuberculosis*"

### 1 Supplementary figures

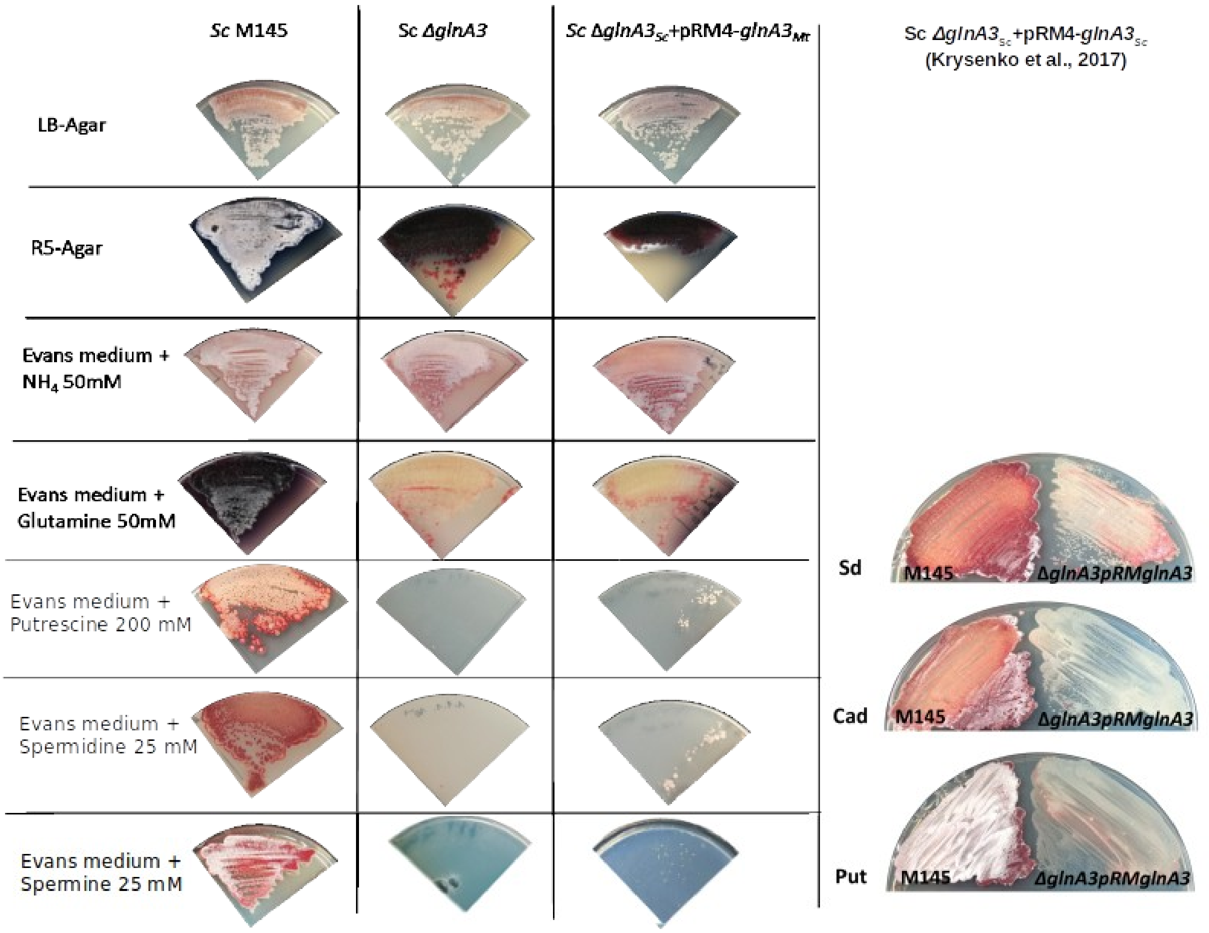


Suppl. Fig. 1. Physiological role of the $glnA3_{Mt}$ gene product in the *S. coelicolor glnA3* mutant grown in the presence of polyamines and other nitrogen sources. Phenotypic comparison of the parental strain *S. coelicolor* M145, the *glnA3* mutant and the *ΔglnA3Sc*pRM4$glnA3_{Mt}$ complementation strain grown on rich LB-agar, R5-agar or defined Evans medium supplemented with glutamine (50 mM), $NH_4Cl$ – ammonium chloride (50 mM) and polyamines putrescine (200 mM), spermidine or spermine (25 mM) as sole nitrogen source. Each panel represents observations on a single agar plate. Complementation of the *S. coelicolor glnA3* mutant with $glnA3_{Mt}$ resulted in restored growth of the strain in the presence of high putrescine, spermidine and spermine concentrations.

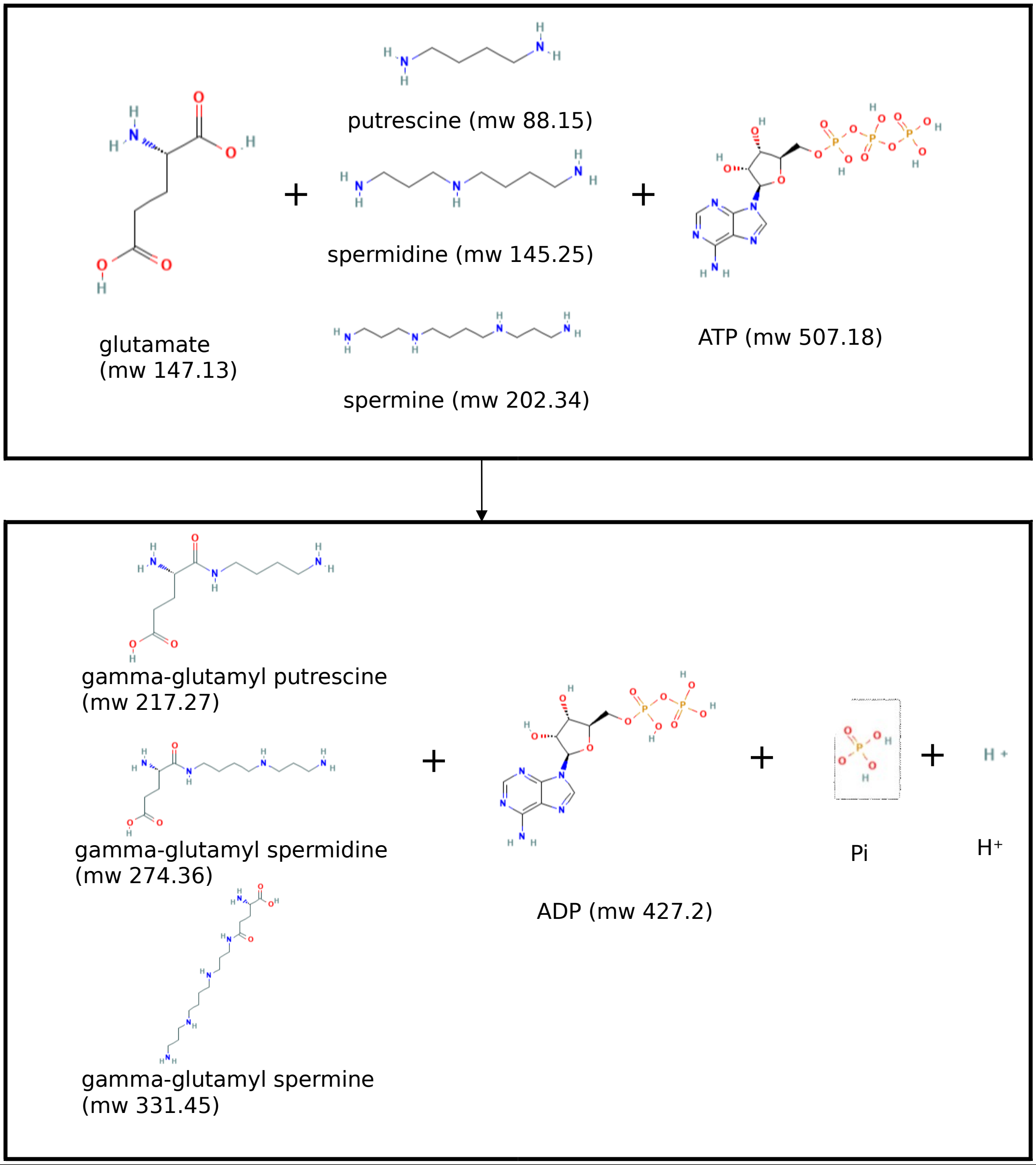


Suppl. Fig. 2. Reaction catalyzed by gamma-glutamylpolyamine synthetase with indicated substrates, products and molecular weight of each molecule.

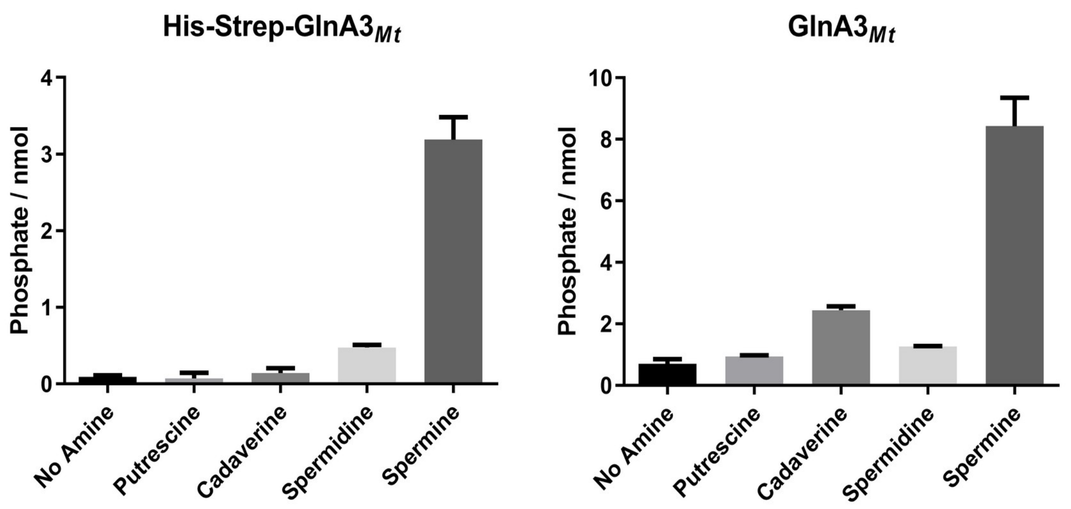


Suppl. Fig. 3. Specific activity of the native GlnA3$_{Mt}$ with different polyamines. All substrates are at a concentration of 50 mM. The mean value of minimum n=3 biological replicates with n=3 technical replicates each with standard error is shown.

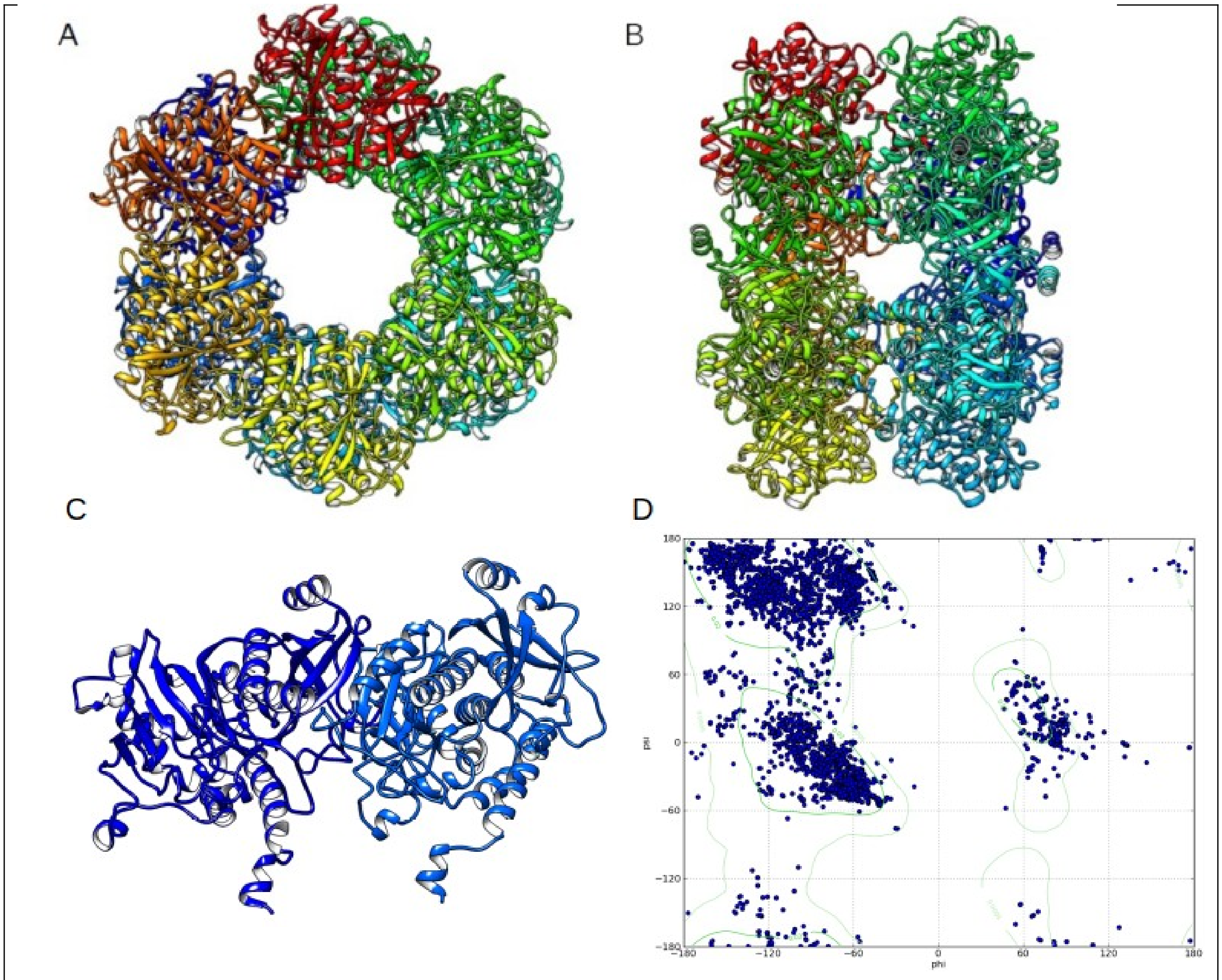


C

Suppl. Fig. 4. Structural alignment of the 3D model structure of GlnA3$_{Mt}$, based on and superposed with the GlnA$_{Mt}$ template (PDB code 1BVC (27)). A) each of the twelve subunits is marked in a different color (color scheme "chain"), B) side-view on two 6-unit containing rings, C) a representation of the active site and substrate binding pocket, D) Ramachandran plot demonstrating the good quality of the generated GlnA3$_{Mt}$ model, based on visualization of energetically allowed regions for backbone dihedral angles ψ against φ of amino acid residues in the structure.

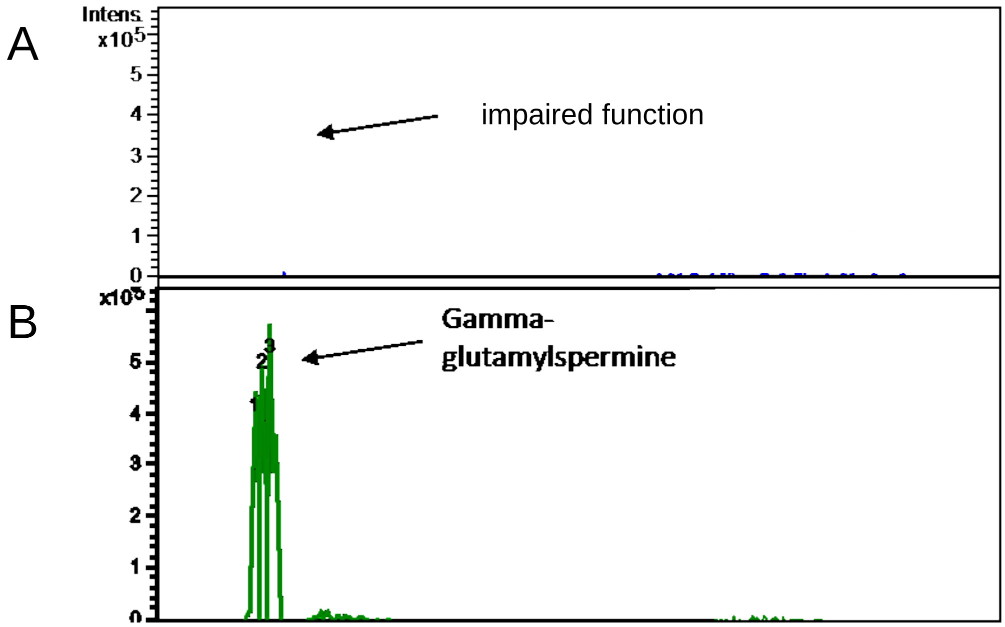


Suppl. Fig. 5. HPLC/ESI-MS analysis of His-Strep-GlnA3$_{Mt}$SER199 and His-Strep-GlnA3$_{Mt}$. Two samples were analyzed in MS negative mode: reaction mixtures with addition of His-Strep-GlnA3$_{Mt}$SER199 (A) and with addition of His-Strep-GlnA3$_{Mt}$ (B). Extracted ion chromatograms for the His-Strep-GlnA3$_{Mt}$ reaction product corresponding to gamma-glutamylspermine with charge to mass ratio of *m/z* 331 was shown (B), and no product in the sample with GlnA3$_{Mt}$SER199 was detected (A).

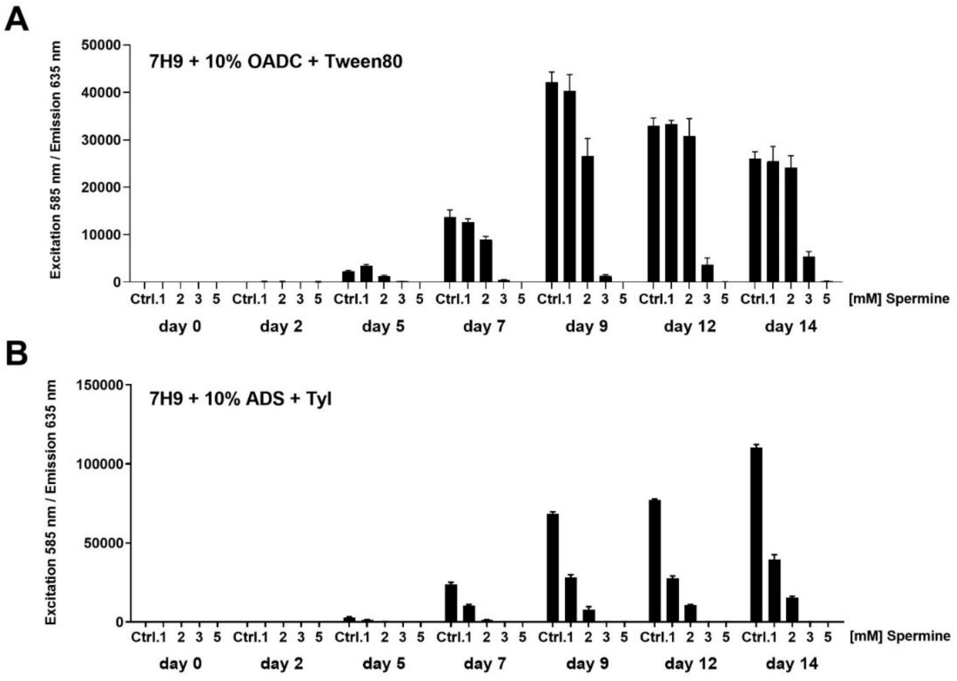

Suppl. Fig. 6. Growth of *M. tuberculosis* in the presence of spermine. *m*Cherry10- expressing Mtb H37Rv bacteria (55) were incubated for a period of 14 days in the absence or presence of spermine in (A) 7H9 medium + OADC + Tween80 or (B) in 7H9 medium + ADS + Tyloxapol and analyzed as described (55); Ctrl: DMSO; Spm: spermine. Concentrations tested were from 1mM to 5 mM.

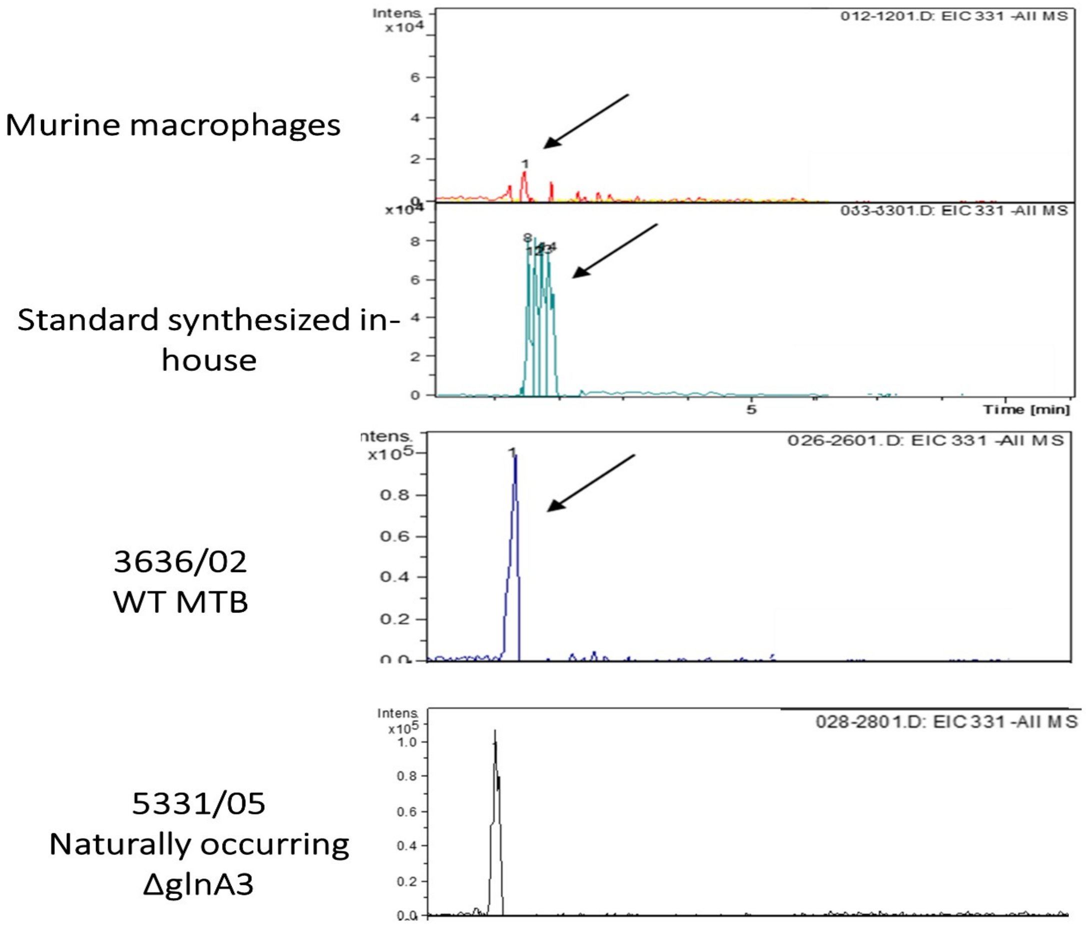


Suppl. Fig. 7. HPLC/ESI-MS detection of gamma-glutamylspermine in the MTB Beijing strain (WT variant) and in the naturally occurring *glnA3* Beijing mutant. Samples were analyzed in MS negative mode.

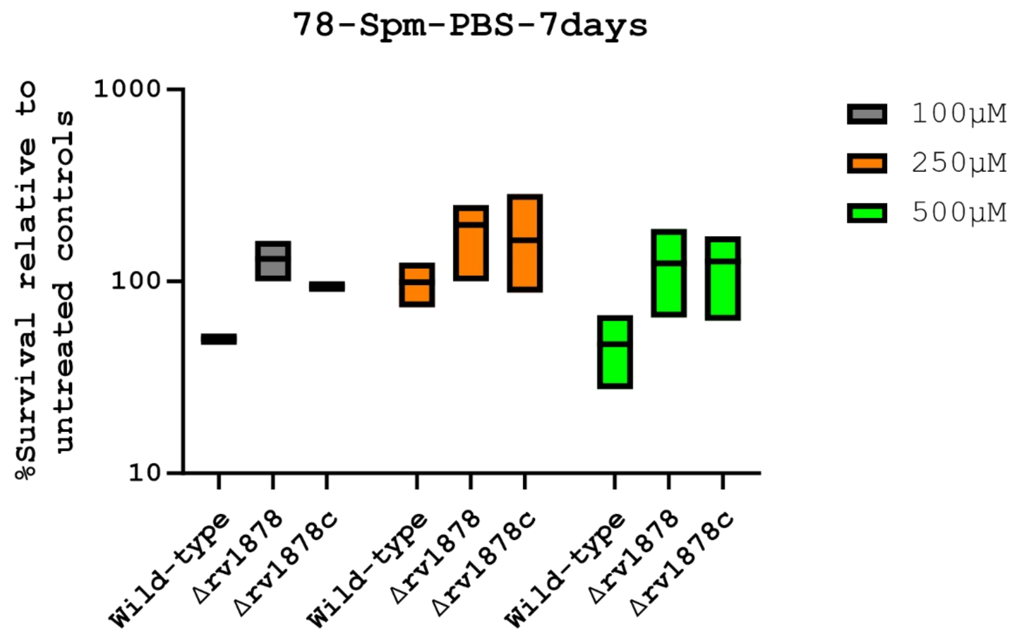


Suppl. Fig. 8. Spermine enhances the survival of the Δ*glnA3* mutant as a sole C/N source.Logarithmic phase cultures (in Sauton's media), were washed several times in PBS, diluted to approximately $10^5$ CFUs/ml, and incubated in the presence (treated) or absence (untreated) of various concentrations of spermine over 7 days. Survival was determined by dividing the number of colonies obtained (after plating strains from various conditions) from the treated conditions to the untreated condition.
